# Microtubule-associated proteins promote microtubule generation in the absence of γ-tubulin in human colon cancer cells

**DOI:** 10.1101/2021.08.13.456214

**Authors:** Kenta Tsuchiya, Gohta Goshima

## Abstract

γ-Tubulin complex acts as the predominant microtubule (MT) nucleator that initiates MT formation and is therefore an essential factor for cell proliferation. Nonetheless, cellular MTs are formed after experimental depletion of the γ-tubulin complex, suggesting that cells possess other factors that drive MT nucleation. Here, by combining gene knockout, auxin-inducible degron, RNA interference, MT depolymerisation/regrowth assay, and live microscopy, we identified four microtubule-associated proteins (MAPs), ch-TOG, CLASP1, CAMSAPs, and TPX2, which are involved in γ-tubulin-independent MT generation in human colon cancer cells. In the mitotic MT regrowth assay, nucleated MTs organised non-centriolar MT organising centres (ncMTOCs) in the absence of γ-tubulin. Depletion of CLASP1 or TPX2 substantially delayed ncMTOC formation, suggesting that they promote MT nucleation in the absence of γ-tubulin. In contrast, depletion of CAMSAPs or ch-TOG did not affect the timing of ncMTOC appearance. CLASP1 also accelerates γ-tubulin-independent MT regrowth during interphase. Thus, MT generation can be promoted by MAPs without the γ-tubulin template.

## Introduction

Microtubules (MTs) are cytoskeletal filaments essential for various cellular activities, such as chromosome segregation, cell division, cell polarisation, and organelle transport. MTs are formed via the polymerisation of α- and β-tubulin heterodimers. MT formation begins with MT nucleation, where tubulin dimers assemble into oligomers and form a ‘critical nucleus’ (Roostalu and Surrey, 2017). The MT nucleus then recruits more tubulin dimers, leading to persistent MT polymerisation or growth, until the MTs pause or start depolymerisation. This entire reaction can take place solely with a high concentration of pure tubulin with GTP in the test tube. However, the initial nucleation step is assumed to be a challenging process *in vivo*, where the tubulin amount is limited and some factors destabilise MTs. The discovery of another class of tubulin, γ-tubulin, and its associated subunits called GCPs, provided key insights into how eukaryotic cells efficiently nucleate MTs (Liu et al., 2021; Oakley and Oakley, 1989; Tovey and Conduit, 2018). The ring-shaped γ-tubulin complex (γ-TuRC) serves as the structural template for the initial tubulin assembly, thereby accelerating the initial lag phase (Zheng et al., 1995). However, the γ-TuRC alone is not an efficient MT nucleator, and efficient nucleation requires association with other proteins, such as CDK5RAP2, XMAP215/ch-TOG, TPX2, and augmin (Alfaro-Aco et al., 2020; Choi et al., 2010; Consolati et al., 2020; Flor-Parra et al., 2018; Tariq et al., 2020; Thawani et al., 2018). Some of these activators likely alter the conformation of γ-TuRC to better fit the ends of MT protofilaments (Consolati et al., 2020; Liu et al., 2020; Wieczorek et al., 2020), whereas others cooperate with γ-TuRC (Consolati et al., 2020; Flor-Parra et al., 2018; King et al., 2020; Thawani et al., 2018). It is now well established that γ-TuRC, with its activators, is the dominant MT nucleator in most eukaryotic cell types. However, there remains another enigma regarding γ-TuRC: cellular MTs are still present after γ-TuRC depletion or perturbation in every system examined to date, including inhibitor treatment and RNA interference (RNAi) in animals and plants (Chinen et al., 2015; Hannak et al., 2002; Nakaoka et al., 2015; Rogers et al., 2008; Sallee et al., 2018; Wang et al., 2015). For instance, RNAi or tissue-specific degradation system reportedly depleted >90% of γ-TuRC from *C. elegans* cells, yet MTs were still nucleated from the centrosome in the early emrbyo (Hannak et al., 2002) or acentrosomal MTOCs in intestinal epithelial cells (Sallee et al., 2018).

This phenomenon is possibly due to the sufficient amount of residual γ-TuRC for a certain degree of MT nucleation. This is not an ignorable caveat. Recent reconstitution studies indicate that a partial complex with eight γ-tubulin subunits is as potent as the full complex of 14 γ-tubulin in facilitating MT nucleation (Wieczorek et al., 2021). To establish the dispensability of γ-tubulin, the best approach is to genetically delete γ-tubulin. However, because γ-tubulin is an essential gene for mitosis in every cell type, it has been impossible to establish a stable cell line in which γ-tubulin genes are deleted. In one study, CRISPR-based genome editing transiently created γ-tubulin gene-deleted cells, which failed to assemble functional spindles (McKinley and Cheeseman, 2017); however, the amount of residual γ-tubulin proteins in each cell was unclear. Another possible explanation for the remaining MTs after γ-tubulin depletion or inhibition is that cells have other factors that can nucleate MTs independent of γ-tubulin. Indeed, several MT-associated proteins (MAPs), whose major activity may not be considered MT nucleation, can promote MT nucleation *in vitro* when mixed with tubulin (Brunet et al., 2004; Imasaki et al., 2021; King et al., 2020; Roostalu et al., 2015; Slep and Vale, 2007). They are candidates for γ-tubulin-independent nucleators in cells.

The aim of this study was to identify the proteins required for γ-tubulin-independent MT nucleation in a single cell type in humans. We first verified that MTs can be nucleated in cells with undetectable levels of γ-tubulin and then searched for the MAPs required for MT generation under these conditions. Our study suggests that multiple factors, including CLASP1 and TPX2, are cellular MT nucleators that are normally masked by the dominant γ-TuRC machinery.

## Results

### MT formation with undetectable levels of γ-tubulin

Most previous studies utilised population assays to assess the contribution of γ-tubulin to MT nucleation, which did not correlate the reduction level of γ-tubulin with MT nucleation potential at the single-cell level. In this study, one of the two γ-tubulin genes in humans (*TUBG2*) was knocked out, and the other TubG1 protein was tagged biallelically with mini-auxin-inducible degron (mAID)-mClover (Fig. 1A, S1A, B). mClover intensity indicated the total γ-tubulin protein level in the cell, whereas mAID allowed acute degradation of the tagged protein via the proteasome. We selected the human HCT116 cell line for this study, which is a stable diploid line derived from colon cancer (Brattain et al., 1981). This cell line is amenable to CRISPR/Cas9-based genome editing and RNAi, and its mitosis has been studied in our laboratory (Okumura et al., 2018; Tsuchiya et al., 2021; Tungadi et al., 2017).

**Figure 1.**
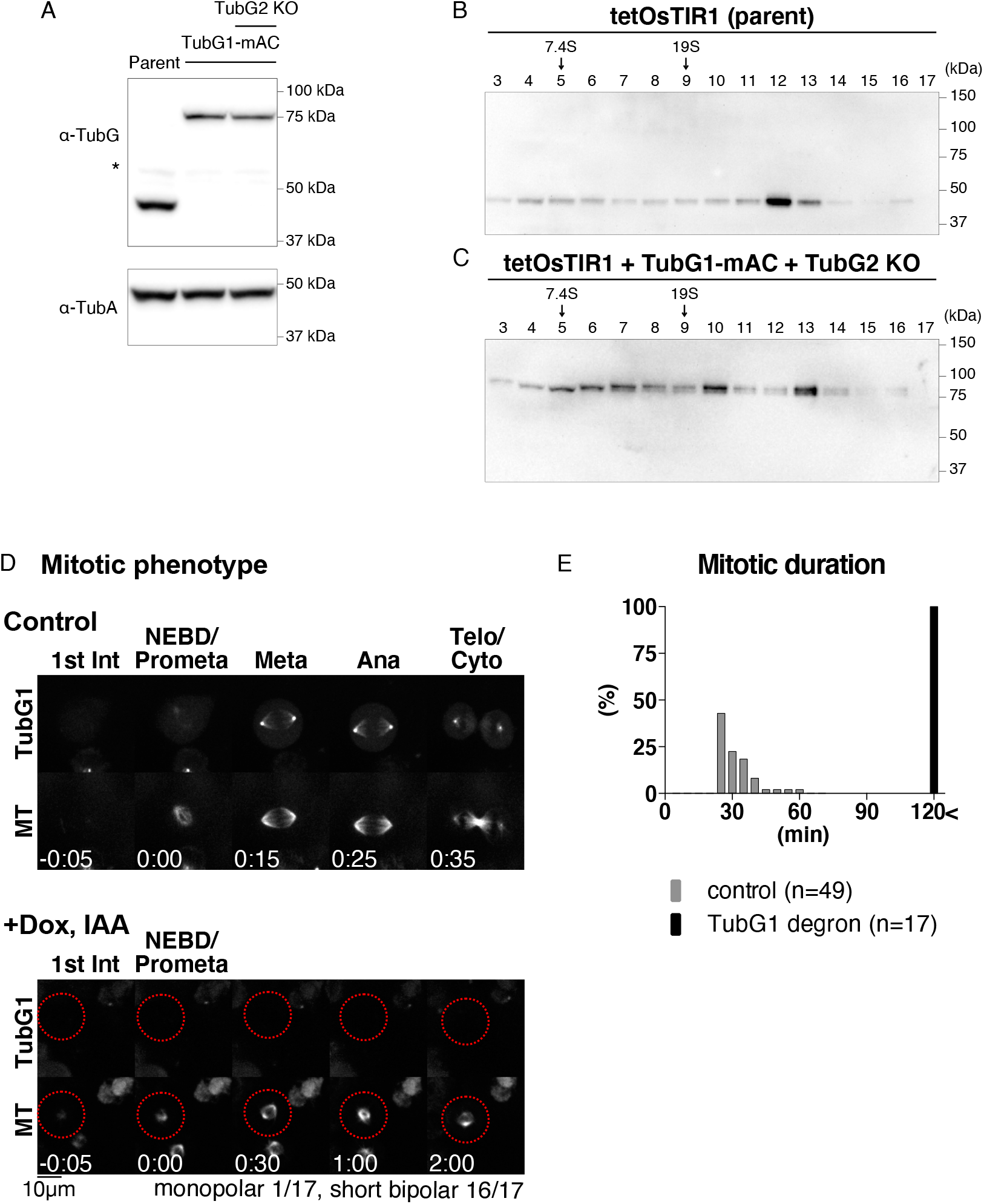
Basic characterisation of TubGl mAID-mClover cell line. (A) Immunoblotting of γ-tubulin and α-tubulin for TubG1-mAID-mClover lines (TubG2 intact and KO lines) and the parental line. (B, C) Sucrose gradient centrifugation followed by immunoblotting of γ-tubulin for parent line (B) and TubG1-mAID-mClover line (C). (D) Spindle dynamics in TubG1-depleted cells (TubG2 KO background) and control cells. Time 0 corresponds to NEBD. (E) Mitotic duration (NEBD to anaphase onset) of γ-tubulin-depleted cells.

Prior to studying MT nucleation, we performed a basic characterisation of this cell line. First, TubG1-mAID-mClover was localised to the centrosome and spindle MTs, consistent with immunostaining in various human cell lines (Luders et al., 2006) (Fig. 1D, Movie 1). Second, the mitotic progression (31 ± 8 min [±SD], n = 49; Fig. 1E) was comparable to that in the control cell line (34 ± 8 min; Tsuchiya et al., 2021). Finally, we performed sucrose gradient centrifugation, followed by immunoblotting (Fig. 1B, C). The results indicated that TubG1-mAID-mClover was assembled into the large γ-TuRC complex. Given the addition of 14 copies of mAID-mClover tag (each ~30 kD), it was reasonable that the peak of the tagged protein was shifted by one lane to a larger fraction than endogenous TubG1. A slightly smaller complex was also detected for tagged TubG1 (lane 10 in Fig. 1C); this may be because the tag partially prevents incorporation into the complete γ-TuRC. Nevertheless, the result is consistent with the observation that endogenous TubG1 can be completely replaced with TubG1-mAID-mClover in this human cell line.

Upon indole acetic acid (IAA) treatment, cells showed different levels of mClover signals (as observed using a spinning-disc confocal microscope) owing to varying degradation levels in interphase and mitosis (Fig. 2A–C). This was confirmed by the quantification of the mClover signal intensity (Fig. 2D, E). The cells in which we could not detect residual γ-tubulin signals by manual inspection always returned low signal values after quantification (coloured pink in the graph). However, the opposite was not true; the cells with very low signals in quantification did not always represent γ-tubulin-null based on manual inspection; they included cells with faint punctate mClover signals at the centrosome, which did not contribute markedly to the total intensity. Therefore, in the subsequent analysis, we manually inspected the acquired images and selected “mClover signals undetectable” cells; these cells were closest to γ-tubulin null. The neighbouring cells with mClover signals served as internal controls. Regardless of the presence or absence of mClover signals, MTs visualised with SiR-tubulin were present in every cell during interphase (Fig. 2A). Furthermore, the cells assembled tiny spindles in mitosis and could not enter anaphase (Fig. 1D, E, Movie 2). These results support the presence of γ-tubulin–independent MT nucleation during interphase and mitosis.

**Figure 2.**
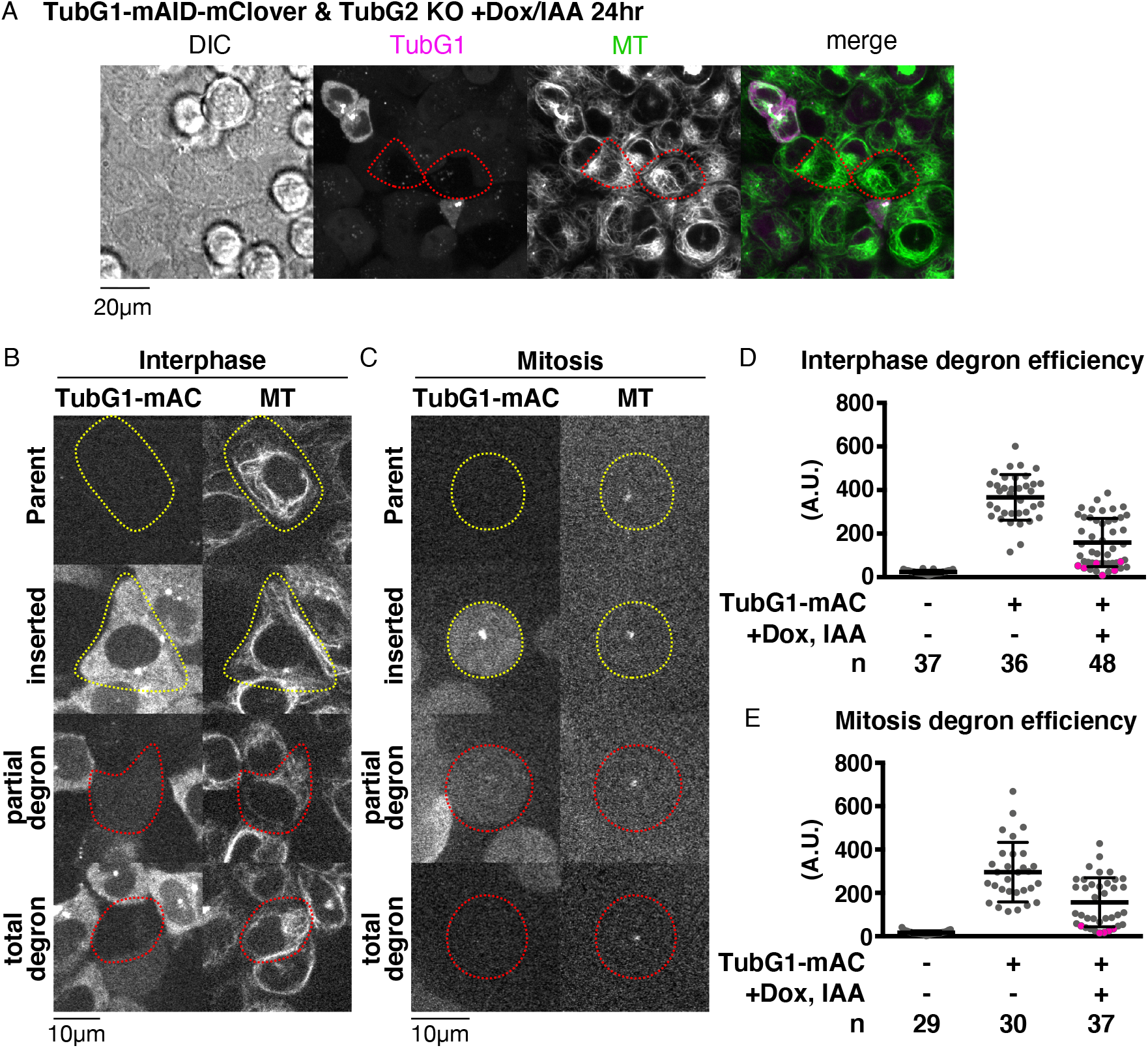
Quantitative assessment of γ-tubulin depletion by AID. (A) MTs are present in cells with undetectable levels of γ-tubulin. Dox/IAA treatment induced degradation of γ-tubulin-mClover in majority of the cells (two representative cells are marked in red circles). (B, C) TubGl-mClover signals in interphase (B) and mitosis (C). Cells circled in red or yellow circles were treated or untreated with Dox/ IAA, respectively. Cells in mitosis were treated with nocodazole to depolymerise most MTs (remaining punctate signals correspond to centrioles). (D, E) Quantification of the mClover signal intensity in the indicated cell lines in interphase (D) and mitosis (E). Total cellular signal intensity was measured at a single focal plane that contained centrioles. Magenta-coloured dots indicate the cells for which an observer manually judged as “signals undetected”.

### MT nucleation in the absence of γ-tubulin

To assess MT generation ability without γ-tubulin in living cells, we performed an MT depolymerisation/regrowth assay, in which MTs were first depolymerised with the MT drug nocodazole, followed by drug washout (Fig. 3A, Movie 3). The experiments were mostly performed at 25 °C, as MTs reappeared too quickly after drug washout at 37 °C; MT nucleation took place prior to image acquisition. For normal HCT116 cells, 25 °C is a challenging temperature, as evidenced by the fact that bipolar spindle formation requires > 30 min (Fig. S2A, B). However, this was compensated for in the regrowth assay, as the initial tubulin concentration was higher than that in normal cycling cells due to complete MT depolymerisation beforehand. In cells that retained TubG1-mClover signals (circled yellow), cytoplasmic MTs were observed 10 min after drug washout, with the centrosome being the most prominent MT organising centre (MTOC) (43 out of 45 cells showed cytoplasmic MT network at 10 min) (Fig. 3B). In contrast, cytoplasmic MTs were hardly detected until 20–30 min in the absence of γ-tubulin, with no clear MTOCs (circled red; 33 out of 42 cells showed no cytoplasmic MT networks at 10 min).

**Figure 3.**
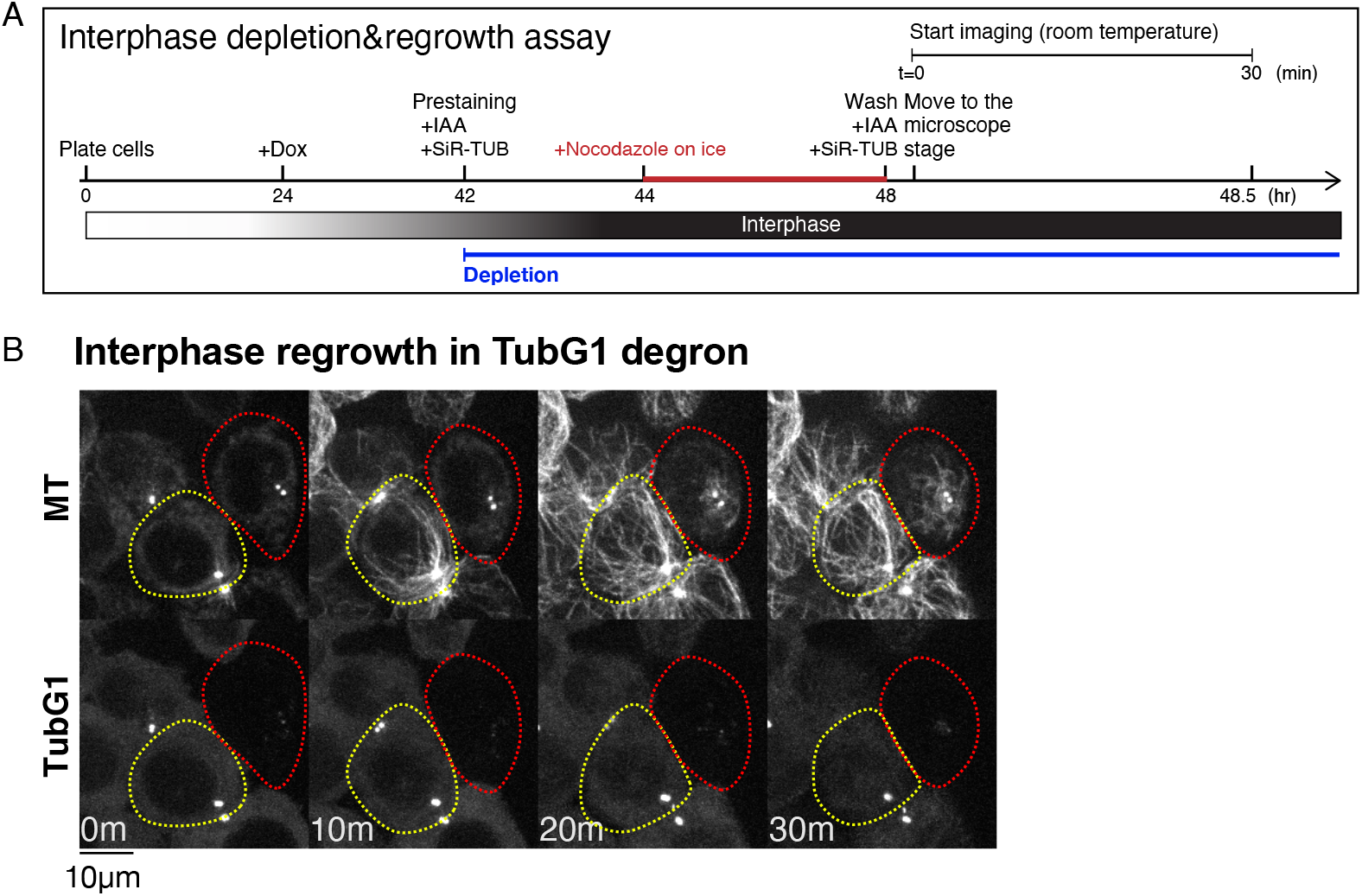
MT nucleation without γ-tubulin. (A) Flowchart of interphase MT depolymerisation-regrowth assay combined with auxin-induced degron. (B) γ-Tubulin-independent MT generation in interphase cells. MTs were depolymerised, except at the centrioles (0 min), followed by induction of regrowth (10–30 min). In the presence of γ-tubulin, MTs start to regrow within 10 min (yellow), whereas it took >10 min in the absence (red). The faint signals after TubGl depletion represent autofluorescence.

To further demonstrate that MT nucleation occurs in the absence of γ-TuRC as the nucleator, we observed the cells undergoing MT regrowth using oblique illumination fluorescence microscopy, which is sensitive enough to detect a single γ-tubulin complex containing >10 fluorescent mClover molecules and an occasional MT nucleation event (Nakaoka et al., 2015). We observed many punctate signals in control cells, each likely representing a cytoplasmic γ-TuRC near the cell cortex (Fig. 4A, yellow circle). We could not identify a MT nucleating event from the observed γ-TuRC spots; MT emergence under these conditions represented MT plus ends grown from the other focal plane. This was because MT nucleation predominantly occurred at the centrosome, which could not be localised to the focal plane in this microscopy. In contrast, after degron treatment, some cells hardly showed punctate signals of mClover, despite the presence of MTs (Fig. 4B, red circle). MTs were generated in the absence of γ-tubulin (i.e. undetectable levels of mClover signals), albeit more slowly (Fig. 4C). Under these conditions, we occasionally identified MT nucleating events, in which MT punctae diffused in 2D, which is an indicator of nucleation rather than plus-end growth from the off-focal plane (Fig. 4D, arrows). Furthermore, we observed at higher frequency the MT loop formation in which both ends were clearly in the focal plane (Fig. 4B, right, Fig. 4D, bottom right, Movie 4); the diameter of the loop was 0.85 ± 0.26 μm (±SD, n = 29), which resembles what has been observed in an *in vitro* MT gliding assay (Liu et al., 2011).

**Figure 4.**
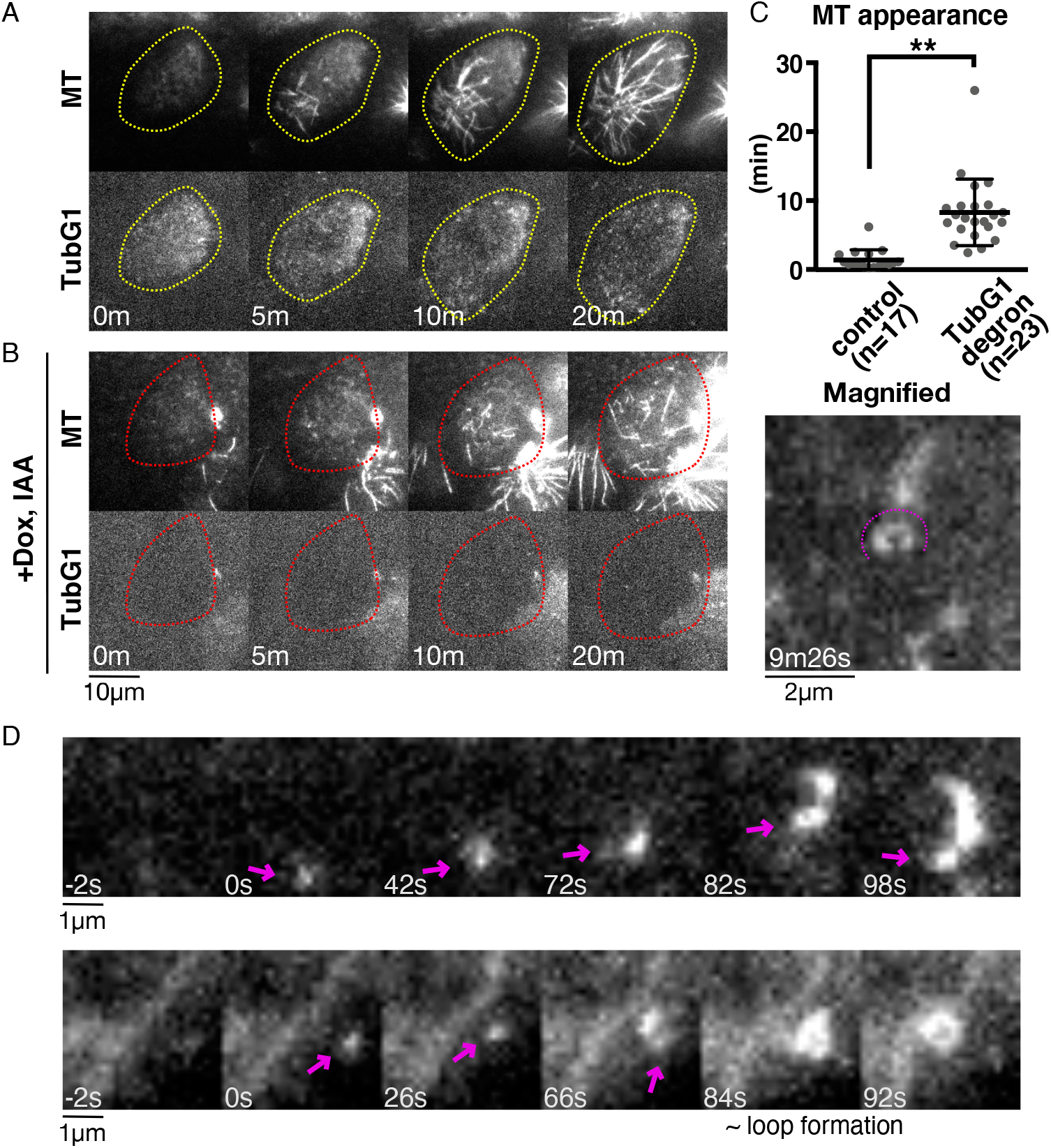
Visualisation of γ-tubulin-independent MT nucleation. (A, B) MT nucleation in the presence (A) or absence (B) of γ-tubulin. Images were acquired every 2 s with oblique illumination fluorescence microscopy, which allows the detection of individual γ-tubulin complex. (C) Time of first MT appearance after drug washout in the presence (1.4 ± 1.5 min [SD]) or absence (8.3 ± 4.8 min) of γ-tubulin. MT was counted when the first SiR-tubulin signal stronger than the background was detected in three consecutive frames (i.e. 6 s). p < 0.0001 (unpaired t-test with Welch’s correction). (D) Two examples of nucleating MTs in the absence of γ-tubulin. The minus ends of nucleating MTs are marked with arrows. MT loop is formed in the second example (84–92 s).

To determine whether there is a possible artifactual effect of SiR-tubulin dye on MT nucleation and growth ability, we compared the timing of MT appearance and MT growth rate in the presence or absence of SiR-tubulin. To visualise MTs without SiR-tubulin, we selected and used a cell line in which endogenous ch-TOG was tagged with mCherry (Fig. S1C, S2C, Movie 1). The data indicate that the effect of SiR-tubulin on MT nucleation and growth is mild (Fig. S2D, E).

Taken together, we concluded that γ-TuRC constitutes the dominant, but not essential, mechanism of MT nucleation in the interphase cytoplasm.

### ch-TOG, CLASP1, and CAMSAPs are critical for interphase MT generation in the absence of γ-tubulin

To identify the factors responsible for γ-tubulin-independent nucleation, we conducted an RNAi screen of 11 candidate genes (or gene family) using the γ-tubulin degron line (Fig. 5A). The MT regrowth assay was carried out, and the cells that retained or lacked γ-tubulin-mClover signals were analysed. In the screening, we identified “MT regrowth” when one or more MTs were detected within 30 min under a spinning-disc confocal microscope. When γ-tubulin was present, MTs were observed normally in all RNAi samples. In contrast, we observed that more than half of the γ-tubulin-degraded cells failed to regrow MTs in 30 min when ch-TOG, CLASP1, or CAMSAP1/2/3 were depleted by RNAi (Fig. 5B–E). In contrast, depletion of CDK5RAP2 or PCNT had no effect on γ-tubulin-independent MT generation, although it has been shown to promote cytoplasmic nucleation in the presence of γ-tubulin in other cell types (Choi et al., 2010; Gavilan et al., 2018; Wu et al., 2016).

**Figure 5.**
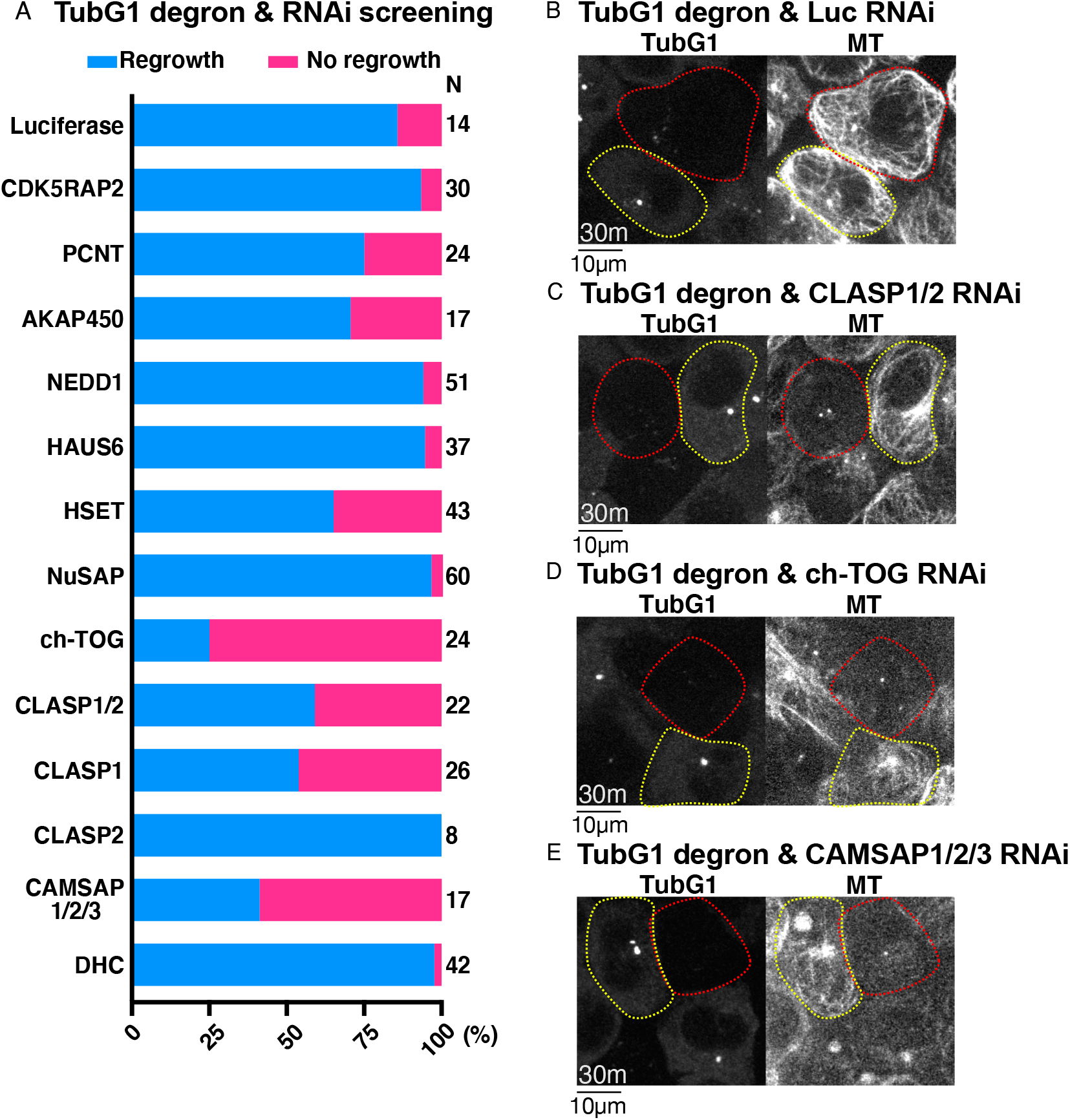
ch-TOG, CLASP1, and CAMSAP1/2 are critical for γ-tubulin-independent MT generation in interphase. (A) Frequency of MT regrowth after RNAi-mediated depletion of the indicated genes (30 min). RNAi cocktails targetted two or three genes simultaneously for CLASP or CAMSAP, respectively. Luiciferase siRNA was used as a negative control. (B-E) γ-Tubulin degron was combined with RNAi of the indicated genes. Images were taken 30 min after nocodazole washout. Cells with γ-tubulin signals are indicated by yellow circles, whereas red-circled cells have no detectable γ-tubulin signals.

To confirm and specify the responsible genes, we performed loss-of-function analyses after generating new cell lines (Fig. 6A–C, Movie 3). The critical contribution of ch-TOG (*Xenopus* XMAP215 orthologue), best known as MT polymerase (Brouhard et al., 2008), was confirmed by degron treatment of the line expressing TubG1-mAID-mClover and ch-TOG-mAID-mCherry (Fig. S1C). When neither signal was observed, MT regrowth was undetectable for 30 min in >70% of the cells (Fig. 6A, D). The requirement of AKAP450 (Gavilan et al., 2018; Rivero et al., 2009; Wu et al., 2016) was excluded from the observation of MT regrowth in their verified knockout (KO) lines (Fig. 6D, S1F, S3A). CLASP proteins are best known as MT stabilisers (Al-Bassam et al., 2010; Moriwaki and Goshima, 2016; Yu et al., 2016) and are also required for Golgi- and γ-tubulin-dependent MT nucleation in RPE1 cells (Efimov et al., 2007). CLASP1 was crucial for MT regrowth, as revealed by the generation of a CLASP1-mAID-mCherry degron line in the background of γ-tubulin degron (Fig. 6B, D, S1D). CAMSAP family members have been characterised as minus-end stabilisers (Goodwin and Vale, 2010; Jiang et al., 2014), in which the CAMSAP3 KO line showed normal regrowth of MTs (Fig. S1G, S3B, C). However, when CAMSAP1 or CAMSAP2 was depleted by RNAi in the CAMSAP3 KO line, MT regrowth was not observed in >25% of the cells (Fig. 6C, D, S3D–G). Quantification of MT intensity at 30 min supported these findings (Fig. 6E). These results indicate that ch-TOG, CLASP1, and CAMSAPs are involved in MT generation during the interphase of γ-tubulin-depleted cells.

**Figure 6.**
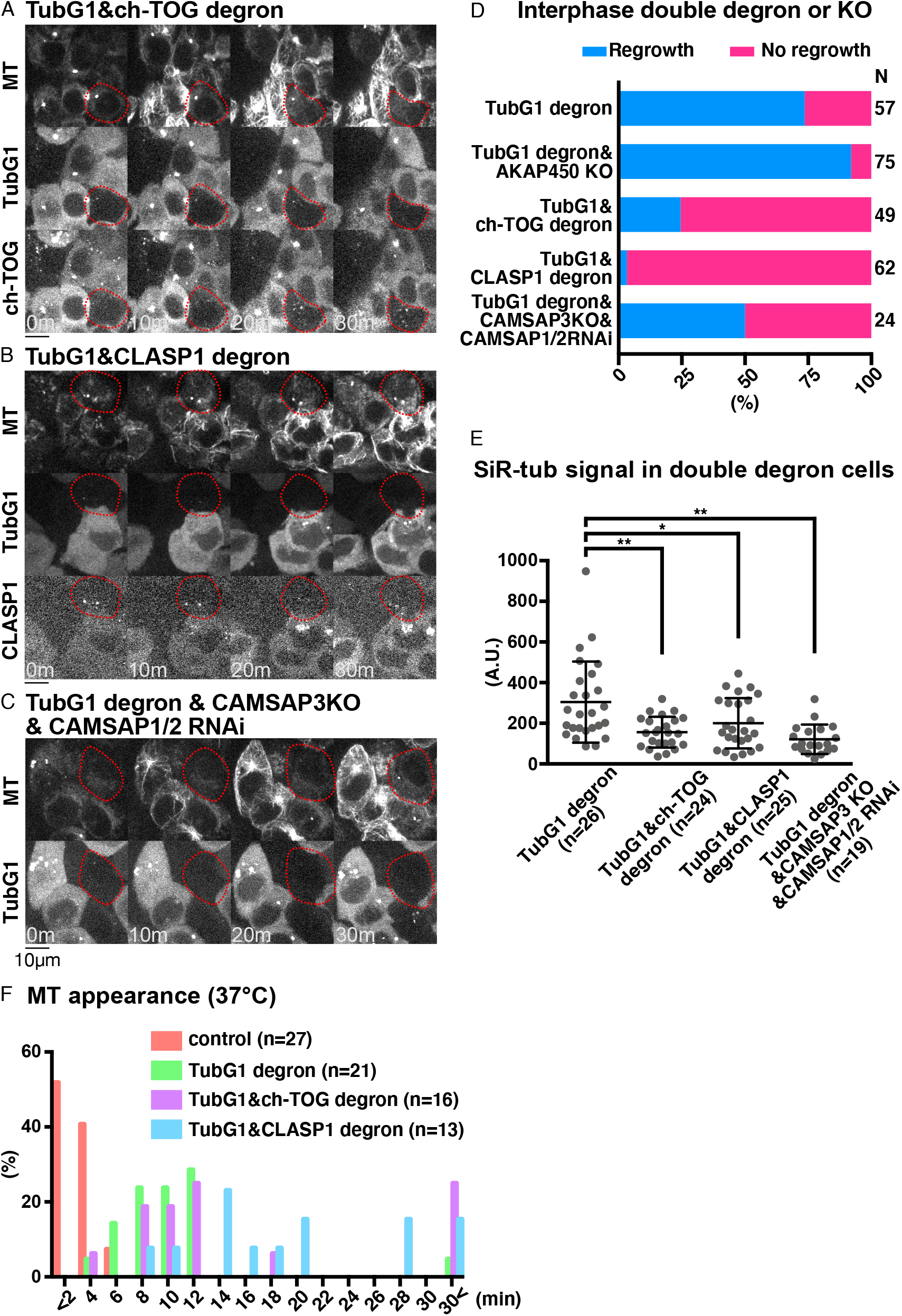
CLASP1 promotes γ-tubulin-independent MT generation. (A-C) Suppression of MT regrowth by double depletion of γ-tubulin and ch-TOG (A), CLASP1 (B), or CAMSAPl/2 (C, CAMSAP3 KO background) at 25°C. Depleted cells are marked in red circles, whereas the surrounding cells with γ-tubulin and MAPs acted as the internal controls. Bars, 10 μm. (D, E) Frequency of MT appearance (D) and MT intensity (E) in the indicated lines (25°C, 30 min after nocodazole washout), p = 0.0009, 0.0313, <0. 0001 (one-way ANOVA, Tukey’s multiple comparisons). (F) Time of MT apperance after nocodazole washout at 37°C in the indicated lines.

An identical assay was performed at 37 °C (Fig. 6F). Even under this more favourable condition for MT nucleation and growth, we observed a significant delay in MT nucleation in the absence of γ-tubulin. Unlike at 25 °C, MTs were observed within 30 min in the majority of cells after co-depletion with ch-TOG or CLASP1. Interestingly, however, the first appearance of MT was delayed when CLASP1, but not ch-TOG, was co-depleted with γ-tubulin (Fig. 6F). These results suggest that CLASP1 is involved in the early stage of MT formation, possibly in the nucleation step, in the absence of γ-tubulin during interphase.

### No accumulation of ch-TOG and CLASP1 at the γ-tubulin-independent nucleation site

We investigated the possibility that ch-TOG or CLASP first forms an assembly or seed, from which MT nucleates and regrows in the absence of γ-tubulin in cells. To this end, the ch-TOG-mCherry or CLASP1-mCherry constructs, which did not have the AID tag, were integrated into the TubG1-mClover-mAID/TubG2-KO line, and oblique illumination fluorescence microscopy was performed after γ-tubulin degradation. We observed MT nucleation and growth after the drug washout. However, ch-TOG or CLASP1 was undetectable at the emergence of MTs; they were later visible near the other end of MTs (Fig. S4). Considering that the SiR-Tubulin dye stains the MT lattice with a ~10 s delay after actual MT formation (David et al., 2019), it is unlikely that ch-TOG or CLASP1 was abundantly present at the nucleation site. We concluded that a detectable level of assembly of ch-TOG or CLASP1 at MT minus ends was not involved in γ-tubulin-independent MT regrowth.

### γ-Tubulin-independent MT nucleation and MTOC formation during prometaphase

Next, we tested the involvement of γ-tubulin in mitotic MT nucleation through MT depolymerisation and regrowth assay in prometaphase (Movie 5). MTs were depolymerised by 24 h nocodazole treatment and 4 h incubation on ice (Fig. 7A). In control cells with γ-tubulin, MTs were undetectable, except for one or two spots (Fig. 7B, 0 min). These most likely reflected centriole-dependent MTs, as MT foci were co-localised with centrin-2 signals in immunostaining images (Fig. S5A), and the punctate signal was not observed when centriole was depleted by a total of 12 days of incubation with centrinone, a chemical inhibitor of the centriole duplication factor Plk4 (Wong et al., 2015) (Fig. S5B). Upon nocodazole washout and returning the cells to 25 °C, MTs were immediately and predominantly nucleated from the centrosomes (Fig. 7B). In the cells in which γ-tubulin was undetectable, MTs similarly disappeared from the mitotic cytoplasm, retaining one or two punctate signals at the centriole (Fig. 7C, 0 min). Upon nocodazole washout, MT regrowth was observed, albeit more slowly, in the absence of γ-tubulin signals (Fig. 7C, 20–30 min). Impaired regrowth was consistent with the results of a previous study in which a γ-TuRC component was depleted in HeLa cells (Luders et al., 2006). Interestingly, in addition to centriolar MTOCs (blue arrow at 6 m 30 s), non-centriolar MTOCs (ncMTOCs) appeared in HCT116 cells, from which MTs later emanated radially (Fig. 7C, right, green arrows). These MTOCs did not have detectable γ-tubulin signals, indicating that MTs are nucleated independent of γ-tubulin or pre-existing MTs. We confirmed that SiR-tubulin staining had negligible impact, as ncMTOCs (visualised by TPX2-mCherry) appeared at similar times and numbers with or without SiR-tubulin staining (Fig. S5C–E). Thus, γ-tubulin is not essential for MT nucleation in prometaphase.

**Figure 7.**
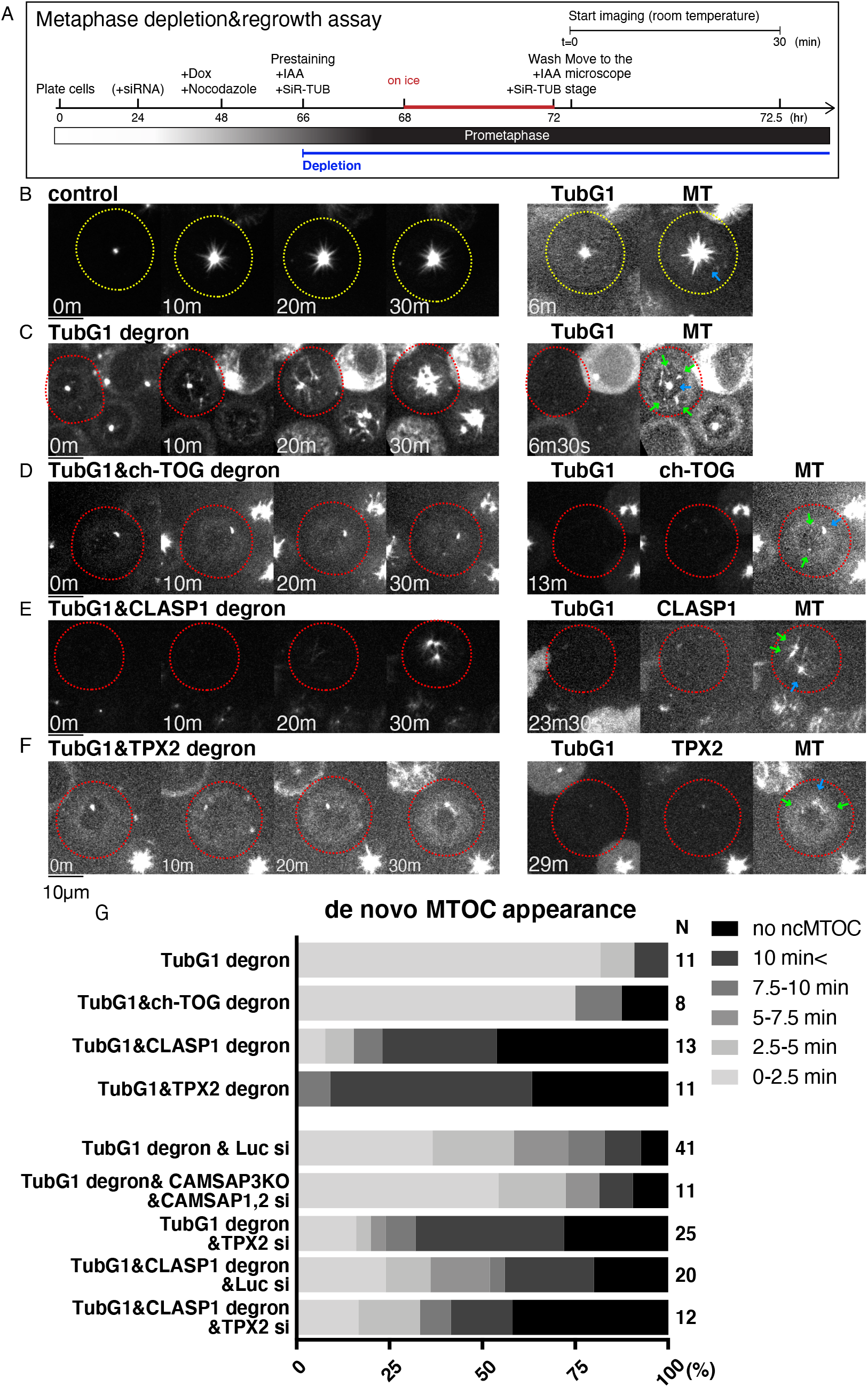
CLASP1 and TPX2 promote γ-tubulin-independent MTOC formation in mitosis. (A) Flowchart of MT regrowth assay in mitosis. (B-F) MT regrowth after drug washout in the indicated cell lines. The cells with undetectable levels of γ-tubulin and MAPs are marked in red circles, whereas the control cells are marked in yellow. Blue arrows on the right panel indicate the centriole, which retains MTs even after cold/drug treatment, whereas green arrows indicate ncMTOCs. Bars, 10 μm. (G) Timing of ncMTOC formation after nocodazole washout in the indicated cell lines.

### TPX2 and CLASP1 promote mitotic MTOC formation, whereas ch-TOG is critical for mitotic MT growth, in the absence of γ-tubulin

ncMTOC is formed through MT nucleation, initial growth, stabilisation, and clustering. We investigated the impact of ch-TOG, CAMSAPs, or CLASP1 depletion on ncMTOC formation in the absence of γ-tubulin.

First, in the absence of ch-TOG alone, ncMTOCs that were clearly separated from centrioles were observed in ~50% of the cells, probably because centrosomal MT growth was suppressed and tubulins were available for other MTOC formation (Fig. S5F, I). ncMTOC was also observed in CLASP1 knockdown; however, centrosomal MTOCs were dominant over ncMTOCs in this case (Fig. S5G, I). In both cases, MT regrowth from MTOCs was not suppressed (Fig. S5J).

Next, we conducted a regrowth assay after co-depletion with γ-tubulin. MT depolymerisation removed most MTs, except one or two punctate centriolar signals, similar to control or single γ-tubulin-depleted cells (Fig. 7D, E, 0 min). In *C. elegans*, co-depletion of γ-tubulin and the ch-TOG orthologue does not result in additional loss of MT regrowth in mitotic cells (Hannak et al., 2002). However, when γ-tubulin and ch-TOG were co-depleted, mitotic MT formation was severely suppressed, consistent with the results of the interphase (Fig. 7D). Interestingly, the initial ncMTOC formation was not substantially affected, indicating that MT nucleation occurred, but the subsequent growth was impaired (Fig. 7D, green arrows, 7G). Similarly, depletion of CAMSAPs did not affect the timing of ncMTOC formation (Fig. 7G). In contrast, after depletion of CLASP1, the appearance of ncMTOCs was dramatically delayed (Fig. 7E, G). These results suggest that ch-TOG is essential for MT growth, but not the initial nucleation step, whereas CLASP1 contributes to ncMTOC formation in the absence of γ-tubulin.

TPX2 plays a role in non-centrosomal MT formation during mitosis in multiple cell lines (Gruss et al., 2002). In the MT regrowth assay using LLC-PK1 and HeLa cells, TPX2 was found to be responsible for non-centriolar MT formation in the presence of γ-tubulin (Cavazza et al., 2016; Katayama et al., 2008; Tulu et al., 2006). We reasoned that this protein might also contribute to ncMTOC formation in the absence of γ-tubulin. We first selected the degron line (Fig. S1E) and performed the mitotic MT depolymerisation/regrowth assay in the presence of γ-tubulin. Similar to the ch-TOG degron, ncMTOCs were observed in ~50% of the cells and MTs regrew from the MTOCs (Fig. S5H–J). This is somewhat different from what has been observed in other studies using different cell lines; in our HCT116 cells, TPX2 was dispensable for ncMTOC formation. To test the contribution of TPX2 in the absence of γ-tubulin, we selected a double-degron line of TPX2 and γ-tubulin, and furthermore, combined TPX2 RNAi with γ-tubulin single or CLASP1/γ-tubulin double degrons. We observed a delay in the appearance of ncMTOCs in either case, indicating that TPX2 promotes ncMTOC formation (Fig. 7F, G). However, ncMTOCs were eventually formed in >50% of the cells in either sample, suggesting that other unknown factors might also nucleate MTs in the absence of γ-tubulin during mitosis.

### Roles of Aurora and Plk1 kinases

Three mitotic kinases have been implicated in mitotic MT generation in previous studies: Plk1/Polo (Cavazza et al., 2016) and Aurora A at the centrosome (Katayama et al., 2008; Magnaghi-Jaulin et al., 2019), and Aurora B in chromosome-proximal regions independent of the centrosome (Carmena et al., 2012). In *C. elegans*, centrosomal MT generation was additively suppressed by depleting γ-tubulin and Aurora A (Motegi et al., 2006). We tested the contribution of these kinases to γ-tubulin-independent MT regrowth in prometaphase by preventing their kinase activity with specific inhibitors in γ-tubulin-depleted cells (Fig. 8A).

**Figure 8.**
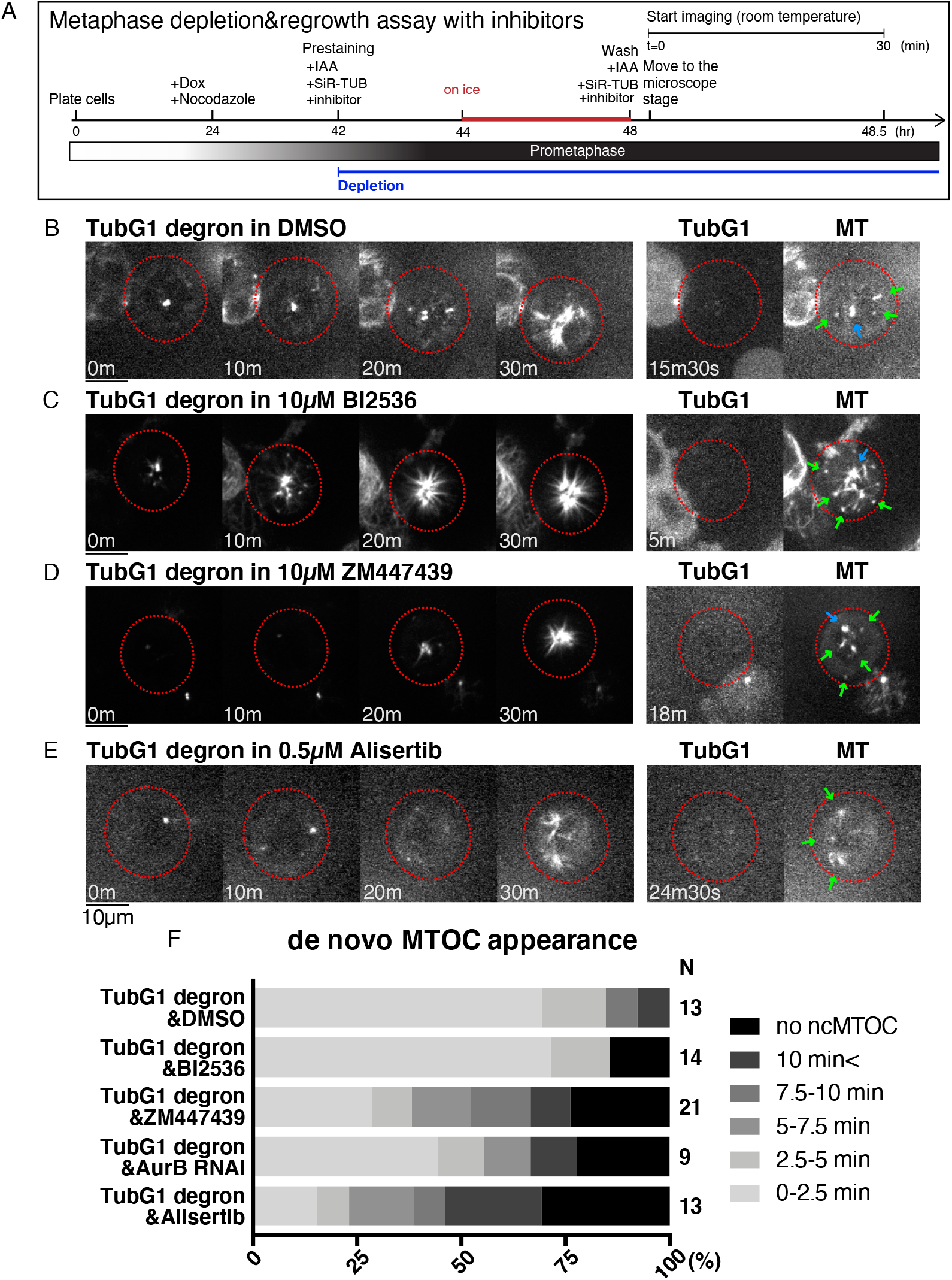
Aurora kinases contribute to γ-tubulin-independent MTOC formation in mitosis. (A) Flowchart of MT depolymerisation-regrowth assay in prometaphase combined with auxin-induced degron and drug treatment. (B–E) MT regrowth after drug washout in the indicated cell lines and treatment. The cells with undetectable levels of γ-tubulin are marked in red circles. Blue arrows on the right panel indicate the centriole, which retains MTs even after cold/drug treatment, whereas green arrows indicate non-centriolar MTOCs. Bars, 10 μm. (F) Timing of ncMTOC formation after nocodazole washout in the indicated cell treatment.

When the Plk1 inhibitor BI2536 was supplied, ncMTOC formed at normal timing, similar to the control γ-tubulin single-depleted cells (Fig. 8B, C, F). In contrast, the inhibition of Aurora B inhibitor by ZM447439 or Aurora A kinase by alisertib delayed ncMTOC formation in the absence of γ-tubulin (Fig. 8D–F). The effect of Aurora B was reproduced by RNAi knockdown of Aurora B. Aurora A and B may be partly involved in non-centriolar MT nucleation in the absence of γ-tubulin.

## Discussion

### MT nucleation in the absence of γ-tubulin

The γ-TuRC is the dominant and arguably the only established cellular MT nucleator in a wide variety of cells. However, experimental perturbation of γ-TuRC in various cell types has never led to a complete loss of cellular MTs. Using a single cell-based assay that monitors both MTs and endogenous γ-tubulin, we unambiguously demonstrated that MTs could be nucleated in the absence of γ-TuRC in human colon cancer cells.

Our functional analysis suggested a possible involvement of MAPs in nucleation. The mitotic MT regrowth assay provided valuable information about non-γ-tubulin MT nucleators, as ncMTOC was visible in almost all cells when γ-tubulin was depleted. The data indicated that TPX2 and CLASP1 contribute to MTOC formation. In *Xenopus* egg extracts and by sophisticated *in vitro* reconstitution, TPX2 was shown to activate augmin- and γ-tubulin-dependent branching nucleation (Alfaro-Aco et al., 2020) and promote template-based nucleation (Wieczorek et al., 2015). Our data suggest that TPX2 not only stimulates the γ-tubulin-dependent process but also potentiates template-free MT nucleation independent of γ-tubulin in the cell. This is consistent with the finding that recombinant TPX2 can promote MT nucleation *in vitro* (Brunet et al., 2004; Gruss et al., 2002; Roostalu et al., 2015). From *in vitro* studies, CLASP1 is best known as a MT stabiliser acting on the MT plus ends and the lattice; it inhibits MT catastrophe and promotes rescue and pausing (Al-Bassam et al., 2010; Moriwaki and Goshima, 2016; Yu et al., 2016). In mitosis, the kinetochore function of CLASP1 has been extensively analysed; however, to our knowledge, the MT nucleation functions have not been extensively discussed (Logarinho et al., 2012; Maffini et al., 2009). CLASPs are known to promote Golgi-mediated MT nucleation during the interphase (Efimov et al., 2007). This function involves AKAP450 and γ-tubulin and is therefore considered to promote γ-tubulin-dependent nucleation (Efimov et al., 2007; Gavilan et al., 2018; Rivero et al., 2009; Wu et al., 2016). It is possible that CLASP1 binds to tubulins and MTs, contributing to the formation of the critical nucleus independent of γ-tubulin. However, because MTOC formation requires not only MT nucleation but also initial growth, stabilisation, and clustering, it cannot be ruled out that TPX2 and CLASP1 regulate the latter three processes.

γ-Tubulin-independent ncMTOC formation was observed in the absence of ch-TOG or CAMSAPs during prometaphase, suggesting that they are not essential for mitotic MT nucleation. MT growth from these ncMTOCs was also observed frequently in the absence of CAMSAPs. In contrast, MT growth from both centriolar and ncMTOCs was inhibited in the absence of ch-TOG and γ-tubulin. We interpret that ch-TOG is dispensable for ncMT nucleation, at least in the presence of CLASP1 and TPX2, but is critical for MT polymerisation, which is consistent with the established role of ch-TOG as the MT polymerase. Regarding centriolar MTOCs, an intriguing possibility is that ch-TOG catalyses centriole-based MT nucleation, independent of γ-tubulin. This is consistent with the proposal on *C. elegans* centrosomes, where the ch-TOG homologue is recruited and concentrates tubulin for nucleation (Woodruff et al., 2017). However, we cannot exclude the possibility that ch-TOG catalyses plus-end polymerisation from nocodazole-resistant MTs at the centriole.

ch-TOG, CAMSAPs, and CLASP1 were important during γ-tubulin-independent MT generation in the interphase. Interestingly, the assay conducted at 37 °C distinguished the phenotype of ch-TOG and CLASP1, which is a favourable condition for tubulin to nucleate and polymerise MTs; MT appearance was delayed specifically in the absence of CLASP1, consistent with the mitosis results. Thus, CLASP1 might be considered involved in the nucleation step, whereas ch-TOG is more critical in MT polymerisation in this cell line. The specific role of CAMSAPs remains unclear, as it is required for MT minus-end stabilisation (Jiang et al., 2014) and might also drive nucleation (Imasaki et al., 2021). In *C. elegans*, the CAMSAP homolog promotes the assembly of non-centrosomal MT arrays in parallel with γ-tubulin (Wang et al., 2015).

Are there other factors redundant with CLASP1 and TPX2 for nucleation? Our protein-depletion experiments did not provide all-or-none results. Most notably, we still observed ncMTOCs in ~30% of the cells after the depletion of γ-tubulin, CLASP1, and TPX2. This might reflect the incomplete depletion of target proteins by AID and RNAi. Alternatively, other MT nucleation factors may also exist. In any case, our data indicate that MT nucleation can be promoted by multiple MAPs in human cells: CLASP1 and TPX2 at the minimum and possibly more.

The mechanism by which CLASP1 or TPX2 promotes nucleation at the molecular level remains unclear. Our imaging suggests that CLASP1 does not form clusters large enough to be visualised by oblique illumination fluorescence microscopy. Thus, the mechanism would be fundamentally different from γ-tubulin-mediated nucleation, where the ring arrangement of 13 γ-tubulin molecules drives nucleation. These MAPs may enhance the longitudinal and lateral contact between tubulins (Roostalu and Surrey, 2017). Regarding TPX2, phase-separated condensates might act as the tubulin concentrator and thereby the MT nucleator (King and Petry, 2020). Furthermore, an interesting *in vitro* study has been published recently, in which the critical nucleus was visualised by electron microscopy (Ayukawa et al., 2021). In their model, the nucleus was characterised by straight tubulin oligomers, which are different from curved tubulin dimers in solution. Provided that this model is correct, TPX2 and CLASP might be considered to convert the curved structure to straight via binding. This is a testable hypothesis *in vitro*.

### Does γ-tubulin-independent nucleation take place in the presence of γ-tubulin?

γ-Tubulin-dependent nucleation at the centrosome is predominant in the HCT116 cell line, and therefore, this study could not determine whether γ-tubulin-independent nucleation occurs in normal HCT116 cells. However, a few reports, besides that of a simple mutant analysis, show that the γ-tubulin-independent mechanism is operating and perhaps important in MT generation in other cell types (Roostalu and Surrey, 2017). One system is the protonemal tissue of the moss *Physcomitrium patens* in which oblique illumination fluorescence microscopy was applied, and the γ-tubulin complex and MTs could be simultaneously observed (Nakaoka et al., 2015). While 90% of the nucleating MTs had γ-tubulin signals at the minus ends, no signals were identified in the other 10% of wild-type cells. The stability of these MTs was explained by the identification of the plant-specific minus-end binding and stabilising protein Spiral2 (Leong et al., 2018). Another notable system is the non-centrosomal fat body cell in *Drosophila*, where γ-tubulin is dispensable for perinuclear MTOC formation, despite being localised at the perinuclear region with the activators (Zheng et al., 2020). In these cells, MTs are generated by CAMSAP and ninein, which recruit ch-TOG. These MTs play critical roles in nuclear positioning. However, single MT or minus ends cannot be specifically visualised in live in this system; it is possible that γ-tubulin also contributes to MT nucleation under normal conditions. Finally, in the electron tomography of the metaphase spindle, MT ends associated with γ-TuRC (ends are closed) and without γ-TuRC (ends are open) were detected (Kamasaki et al., 2013; O’Toole et al., 2003). The open ends represent either plus ends of MTs or the minus ends of MTs nucleated independent of γ-tubulin. Taken together, it can be assumed that the γ-tubulin-independent mechanism operates and plays a role in the activity of at least certain animal and plant cell types.

## Materials and methods

### Plasmid, cell culture, and cell line selection

Plasmids for CRISPR/Cas9-mediated genome editing and auxin-inducible degron were constructed using standard protocols (Natsume et al., 2016; Okumura et al., 2018). The plasmids and sgRNA sequences used in this study are listed in Tables S1 and S2, respectively. In the normal passage, the HCT116 cell line possessing DOX-inducible tet-OsTIR1 was cultured at 37°C with McCoy’s 5A medium (Gibco) supplemented with 10% serum and 1% antibiotics (Natsume et al., 2016). Knock-in and knockout lines were generated by CRISPR/Cas9 genome editing essentially as previously described (Okumura et al., 2018). CRISPR/Cas9 and donor plasmids were co-transfected into the cell lines using Effectene (Qiagen, Venlo, Netherlands). For drug selection, 1 μg/mL puromycin (Wako Pure Chemical Industries, Osaka, Japan), 800 μg/mL G418 (Roche, Basel, Switzerland), 200 μg/mL hygromycin B (Wako Pure Chemical Industries), and 8 μg/mL blasticidin S hydrochloride (Funakoshi Biotech, Tokyo, Japan) were used. Selection medium was replaced with fresh selection medium 4–5 d after starting selection. After 10–14 d, colonies grown on a 10 cm culture dish were washed once with PBS, picked up with a pipette tip under a microscope (EVOS XL, Thermo Fisher Scientific, Waltham, MA) located on a clean bench, and subsequently transferred to a 96-well plate containing 50 μL of trypsin-EDTA. After a few minutes, the trypsinized cells were transferred to a 24-well plate containing 500 μL of the selection medium, and then further transferred to a 96-well plate (200 μL per well) for the preparation of genomic DNA. For the preparation of genomic DNA, cells in the 96-well plate were washed once with PBS and lysed by 90 μL of 50 mM NaOH. After boiling for 10 min, the solution was equilibrated by 10 μL of 100 mM Tris-HCl (pH 9.0). To confirm the genomic insertion, PCR was performed using 1–2 μL of the genomic DNA solution and Tks Gflex DNA polymerase (Takara Bio). The primers are listed in Table S3. For the TubG1-KO/TubG2-mClover-mAID line, γ-tubulin and α-tubulin were immunoblotted with monoclonal antibodies GTU88 (Sigma, 1:10,000) and DM1A (Sigma, 1:2,000), respectively, and the lack of untagged γ-tubulin protein was confirmed. Proper tagging to ch-TOG, CLASP1 and TPX2, and biallelic deletion of CAMSAP3 and AKAP450 were confirmed by immunoblotting with specific antibodies as follows: ch-TOG (QED Bioscience, 1:1,000), CLASP1 (Abcam, 1:1,000), TPX2 (anti-rabbit, 1:200, a gift of Dr. Isabelle Vernos (Gruss et al., 2002)), CAMSAP3 (anti-rabbit, 1:200, gift of Dr. Masatoshi Takeichi (Tanaka et al., 2012)) and AKAP450 (anti-rabbit, 1:1,000, gift of Dr. Yoshitaka Ono (Takahashi et al., 1999)). All tagged lines grew in a manner that was indistinguishable from the parental line, indicating that the tag did not substantially affect protein function. To activate auxin-inducible degradation, cells were treated with 2 μg/mL Dox for 20–24 h and 500 μM IAA for the duration indicated in each figure. RNAi was performed using Lipofectamine RNAiMAX (Invitrogen), following manufacturer’s instruction.

### Biochemistry

Immunoblotting was performed using a standard protocol with SDS sample buffer, except for ch-TOG detection, which might be prone to degradation during this procedure. For ch-TOG, cells were treated with 4M urea-containing sample buffer for 10 min at room temperature (Ito and Goshima, 2015). Sucrose gradient centrifugation was performed according to previously reported methods (Choi et al., 2010; Teixido-Travesa et al., 2010). Confluent cells on three 10-cm culture dishes were lysed with 800 μL lysis buffer (50 mM HEPES-KOH pH 7.6, 150 mM NaCl, 1 mM EGTA, 1 mM MgCl2, 1 mM DTT, 0.5% NP-40, 100 μM GTP, and protease inhibitors), followed by 27 G needle passages.

After two rounds of centrifugation (13,000 rpm, 15 min in a tabletop centrifuge and 50,000 rpm, 15 min in TLA100.3 rotor [Beckmann]), 500 μL supernatant was loaded onto a 10%–40% sucrose gradient in a SETON tube (#7022), which was prepared using Gradient Station (BIOCOMP), and centrifuged in an MLS-50 rotor (Beckmann) at 50,000 rpm for 3 h 45 min at 4 °C. Fractionation was performed with Gradient Station attached to MicroCollector (AC-5700P, ATTO). Aldolase (7.4S) and thyroglobulin (19S) were used as size markers.

### Microscopy

Imaging was mostly performed using spinning-disc confocal microscopy with a 60× 1.40 NA lens (Nikon). A CSU-X1 confocal unit (Yokogawa Electric Corporation) and an EMCCD camera ImagEM (Hamamatsu Photonics) were attached to a Ti-E inverted microscope (Nikon) with a perfect focus system. Several DIC images were acquired with another spinning-disc confocal microscope, in which CSU-W1 and ORCA-Flash4.0 digital CMOS camera (Hamamatsu Photonics) were attached to Ti-E (courtesy of Dr. Tomomi Kiyomitsu). Oblique illumination fluorescence microscopy was performed following the method used for plant cell imaging (Nakaoka et al., 2015). Briefly, the cortical region of interphase cells on the glass-bottom dish was imaged every 2 s with a Nikon Ti-E microscope equipped with an EMCCD camera Evolve (Roper) and the total internal reflection fluorescence unit and a 100 × 1.49 NA lens (Nikon). A fragment of broken glass was placed on the sample to flatten the cells. Imaging for regrowth assay was performed mostly at 25–26 °C and sometimes at 37°C as indicated in the figure. Time-lapse imaging of regular mitosis and the degron efficiency analysis were performed at 37°C. The microscopes were controlled using NIS-Elements software (Nikon). Centrin-2 immunostaining was performed with a specific antibody (SantaCruz, rabbit, 1:500) after methanol fixation. All image analyses of live spinning-discs were based on maximum projection images, whereas a single focal plane was shown for the immunofluorescence image and measurement of SiR-tubulin signals. To optimise the image brightness, the same linear adjustments were applied using Fiji. MT growth in prometaphase was determined at 30 min; the appearance of filamentous signals emanating from MTOCs were the indicator of MT regrowth.

### MT regrowth assay

The flowcharts are shown in the figures. In one mitosis experiment, typically 3–4 analysable cells were obtained; to obtain N ≥ 10, at least three independent experiments were performed. Cells were cultured in 4-well glass-bottomed dishes (CELLview™, #627870; Greiner Bio-One, Kremsmünster, Austria) and maintained in a stage-top incubator (Tokai Hit, Fujinomiya, Japan). 5% CO_2_ was supplied. The heater was not turned on, and the experiment was performed at room temperature (~25°C). MTs were stained with 50 nM SiR-tubulin (Spirochrome) for >1 h prior to image acquisition (Lukinavicius et al., 2014; Okumura et al., 2018). Cells in a 4-well glass-bottom dish were treated with 40 ng/mL nocodazole on ice for 4 h (interphase) or at 37°C for 20–24 h (prometaphase), followed by drug washout by medium exchange twice (1 min each, 700–800 μL, room temperature). This incubation time on ice was set as residual MTs or dead cells were detected by shorter or longer incubation, respectively. The specimen was immediately set under a microscope, and images were acquired. In most experiments, the cells were kept at room temperature (~25°C) to prevent MT nucleation before sample setup and slow down the nucleation step. For regrowth assay at 37°C, cells were washed by 37°C warmed medium and the temperature was contoroled by stage top incubator. Images were acquired every 30 s for 30 min with spinning-disc microscopy equipped with a piezo stage (1 μm × 3 or 5 z-sections). The maximum projection images are displayed in the figures. Cells were treated with mitotic kinase inhibitors for 2 h prior to imaging, and imaging was performed in the presence of drug treatments (BI2336, 10 μM; ZM447439, 10 μM; and Alisertib, 0.5 μM). BI2536 was effective at this concentration in the HCT116-TubG1 degron line, as 16 out of 17 cells showed monopolar spindles at only 30 nM. ZM447439 was shown to be effective in HCT116 cells at concentrations of 2 μM or higher (Dreier et al., 2009; Li et al., 2010). Moreover, we reproduced the phenotype using RNAi. The reported IC50 value of alisertib in HCT116 was 0.032 μM (Manfredi et al., 2011) and 0.04 μM (Davis et al., 2015), which is much lower than our applied concentration (0.5 μM). Other studies have shown that p53 is fully activated at 0.4 μM or above (Marxer et al., 2014) or that apoptosis is observed similarly at 0.1 μM and 1 μM (Pitts et al., 2016).

## Acknowledgments

We would like to thank Tomomi Kiyomitsu for cell lines, reagents, and valuable comments on the manuscript, Marie Nishikawa for the selection of a cell line, Tomoko Nishiyama for help of sucrose gradient centrifugation, and Aoi Takeda for checking several cell lines. This work was funded by JSPS KAKENHI (17H01431), and JSPS and DFG under the Joint Research Projects-LEAD with UKRI to G.G. The authors declare no conflicts of interest.

---

**Movie 1. Mitotic localisation of tagged proteins**

mClover-tagged TubG1 and mCherry-tagged ch-TOG, CLASP1, and TPX2 were imaged with MTs (visualised with SiR-tubulin) using spinning-disc confocal microscopy. Time indicates hours and minutes. Bar, 10 μm.

**Movie 2. Spindle phenotype after γ-tubulin depletion**

Mitotic progression after γ-tubulin depletion. MTs (green) were visualised using SiR-tubulin and imaging was performed using spinning-disc confocal microscopy. Magenta; TubG1-mClover. Time is shown in hours and minutes.

**Movie 3. MT regrowth after depolymerisation in interphase**

Indicated proteins were depleted by AID in cells marked in white circles. MTs were visualised by SiR-tubulin and imaged using spinning-disc confocal microscopy. Time indicates minutes and seconds.

**Movie 4. MT regrowth after depolymerisation in interphase-oblique illumination**

MT regrowth in a γ-tubulin-depleted cell was observed using oblique illumination fluorescence microscopy. The region near the cell cortex was visualised using microscopy. Ring-shaped MTs are indicated by arrows. Time indicates minutes and seconds.

**Movie 5. MT regrowth after depolymerisation in mitosis**

The indicated proteins were depleted by AID in the white-circled cells. MTs were visualised by SiR-tubulin and imaged using spinning-disc confocal microscopy. Multiple ncMTOCs were detected in the γ-tubulin single-depleted cells. Time indicates minutes and seconds.

**Table S1.** Plasmids for homologous recombination

| Name | Purpose | Homology arm |  | Selection |
| --- | --- | --- | --- | --- |
|  |  | N term | C term |  |
| pKT1 | TubG1-mClover-mAID integration | 248 bp | 202 bp | G418 |
| pKT11 | TubG2 knock out | 642 bp | 526 bp | blasticidin |
| pKT35 | ch-TOG-mAID-mCherry integration | 600 bp | 669 bp | hygromycin |
| pKT52 | ch-TOG-mCherry integration | 600 bp | 669 bp | hygromycin |
| pKT90 | CLASP1-mAID-mCherry integration | 574 bp | 632 bp | hygromycin |
| pKT43 | TPX2-mAID-mCherry integration | 201 bp | 254 bp | hygromycin |
| pKT53 | TPX2-mCherry integration | 201 bp | 254 bp | hygromycin |
| pKT55 | AKAP450 knock out | 598 bp | 601 bp | hygromycin |
| pKT58 | CAMSAP3 knock out | 834 bp | 585 bp | hygromycin |

**Table S2.**
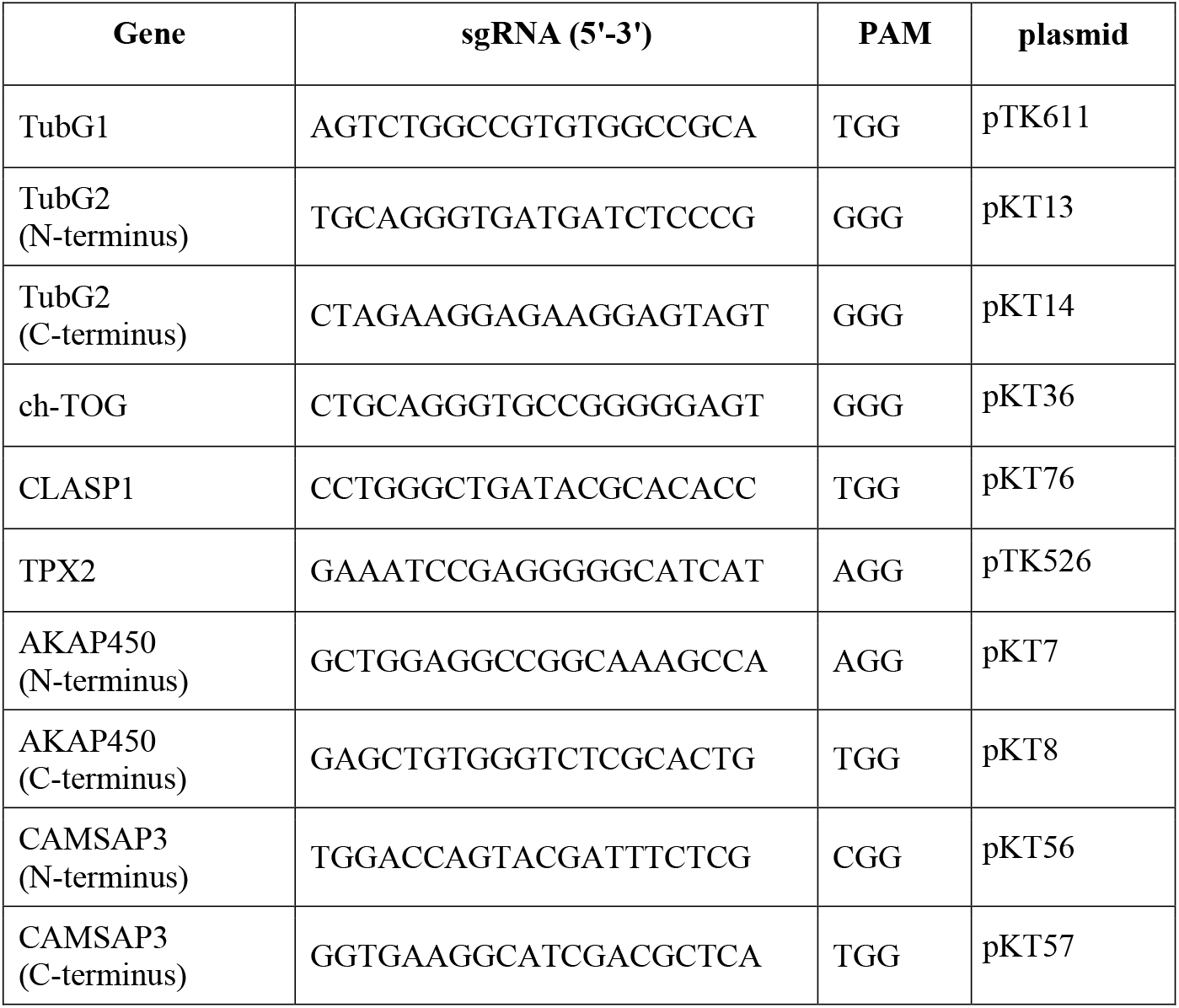
sgRNA sequences for CRISPR/Cas9-mediated genome editing

**Table S3.**
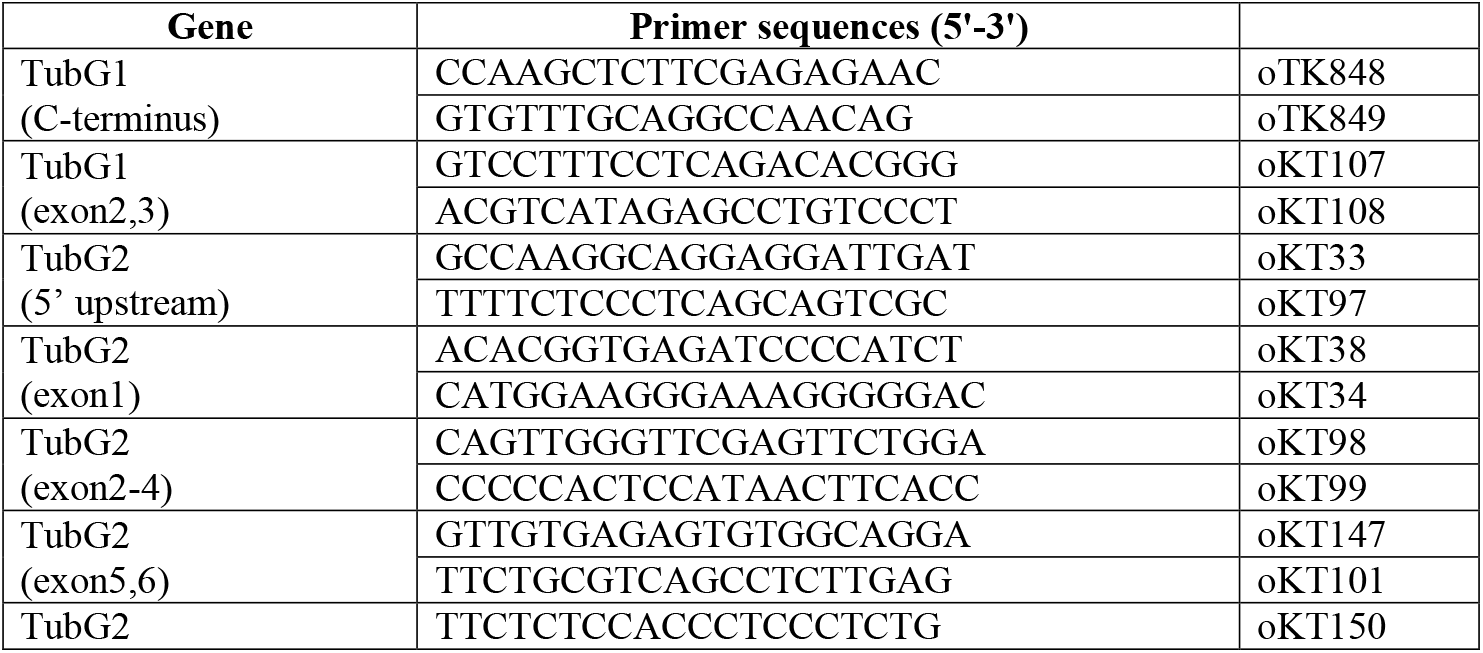
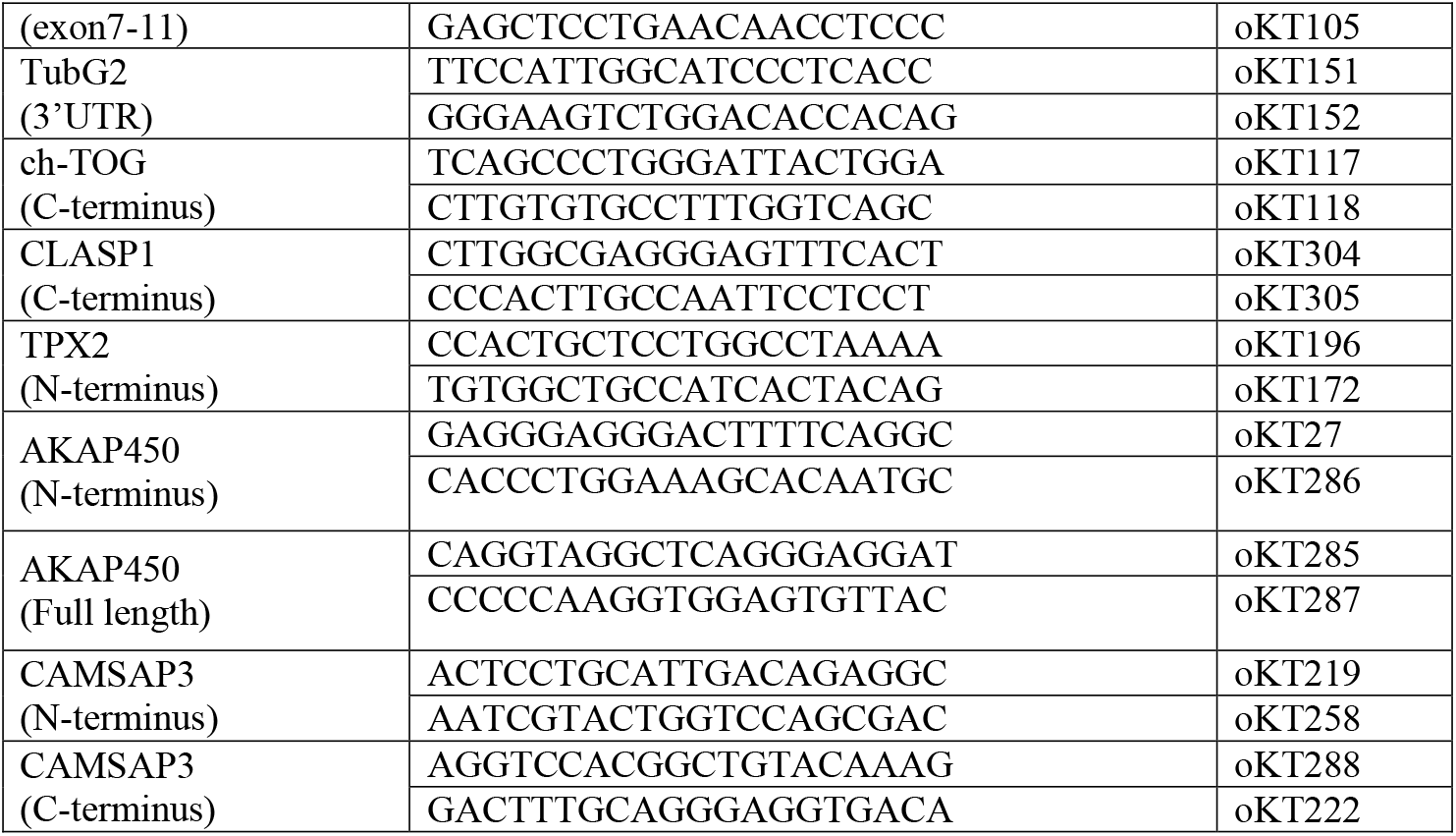
PCR primers to confirm gene editing

**Table S4.**
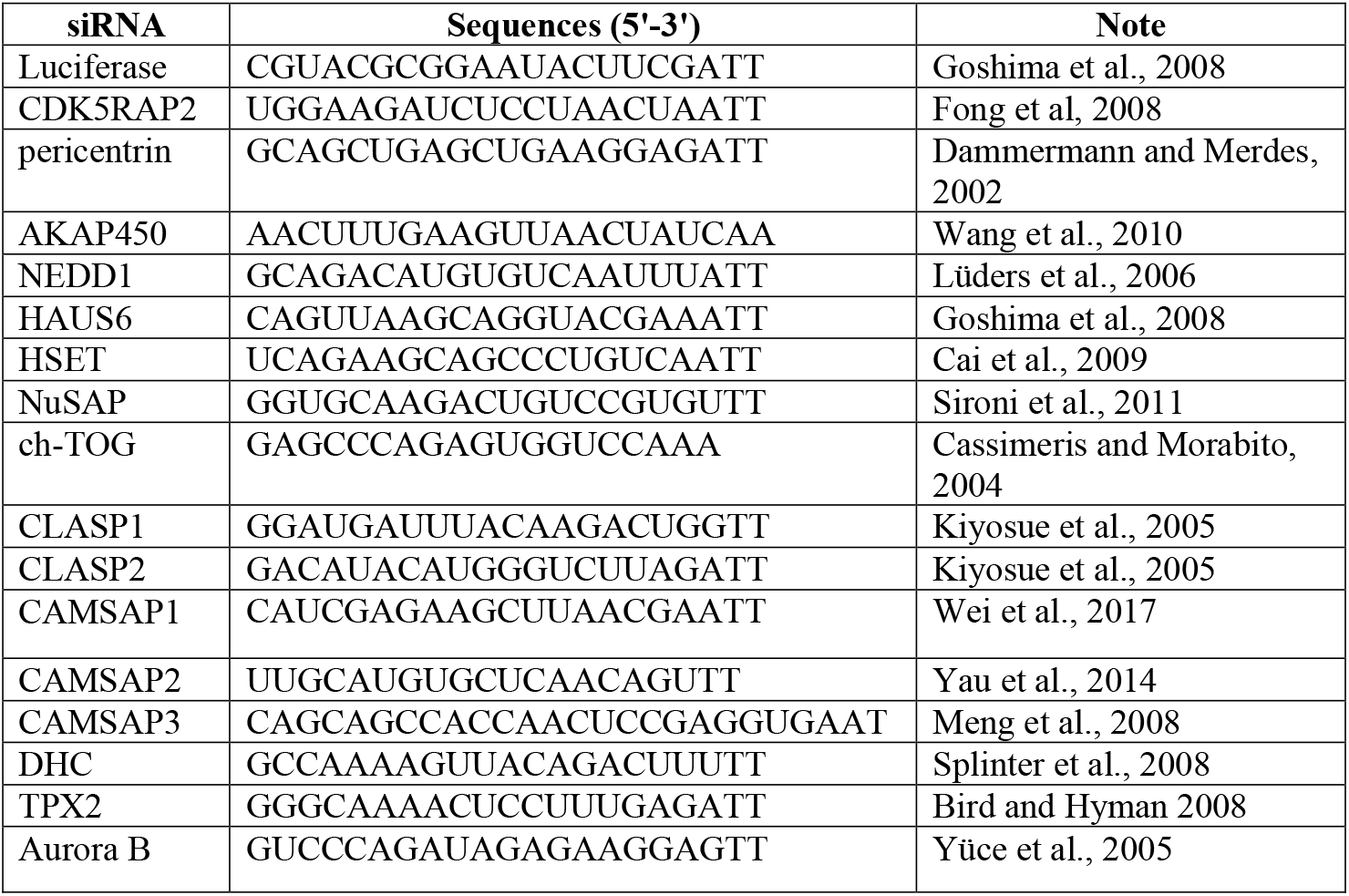
Primers for RNAi

**Figure S1.**
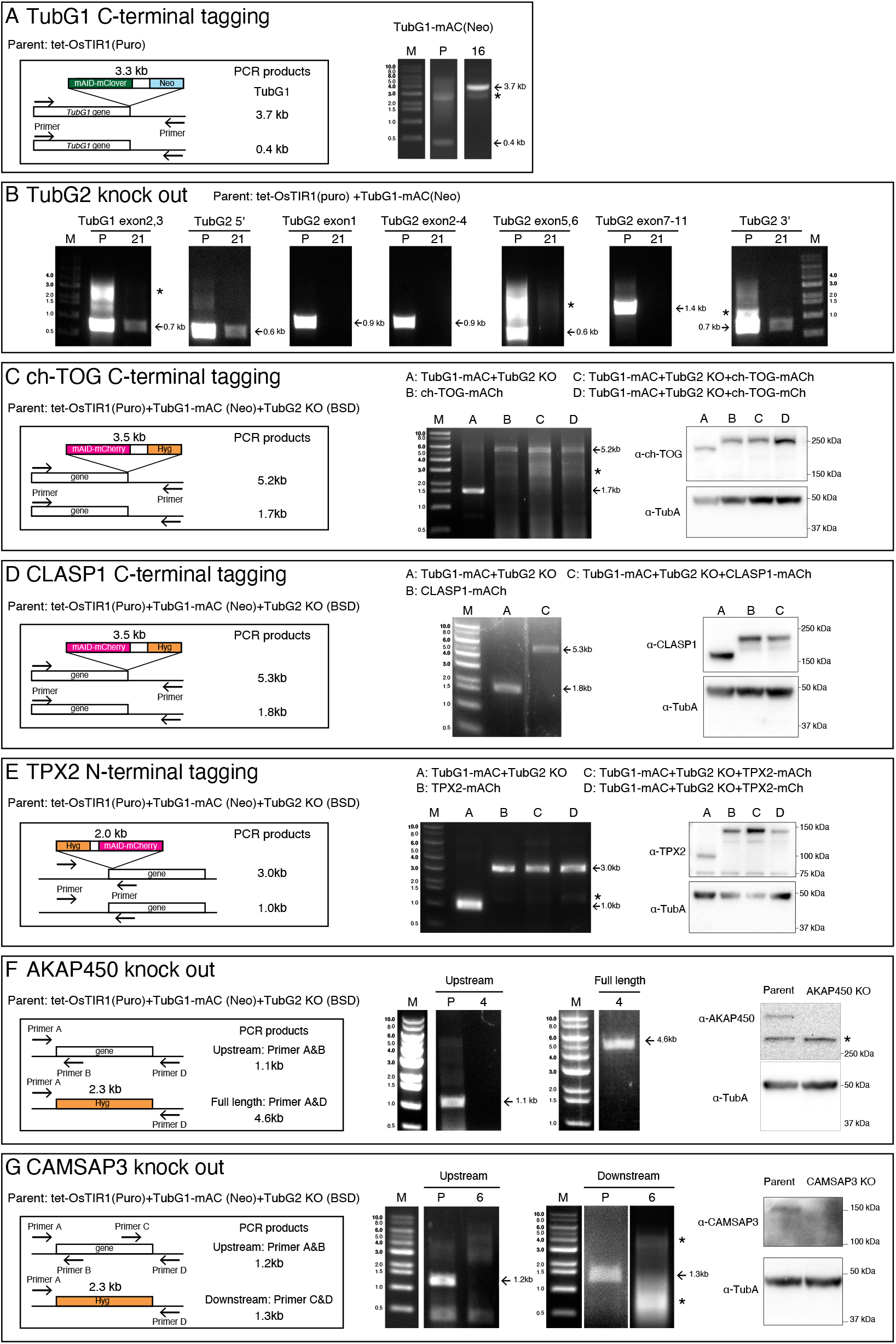
Construction and confirmation of the cell lines used in this study. Gene targeting strategy, PCR strategy, PCR results, and immunoblotting results are shown for the cell lines used in this study. ‘P’ marks at the top of the panels represent parental lines, whereas the numbers (e.g. ‘4’, ‘16’) indicate the line identification numbers. Asterisks indicate non-specific bands. (A) mAID-mClover (‘mAC’) tagging to TubGl. PCR amplified a 3.7-kb fragment in the tagged line. Immunoblotting results are shown in Fig. 1A. (B) Confirmation of TubG2 KO by PCR The lack of bands derived from exons was confirmed in the KO line. The TubGl exon and TubG2 UTR regions were amplified as positive controls. (C) mAID-mCherry (‘mACh’) or mCherry (‘mCh’) tagging to ch-TOG. PCR amplified a 5.2-kb fragment in the tagged line. Immunoblotting results are shown on the right; the band is shifted upwards with the tag. (D) mAID-mCherry (‘mACh’) tagging to CLASP1. PCR amplified a 5.3-kb fragment in the tagged line. Immunoblotting results are shown on the right; the band is shifted upwards with the tag. (E) mAID-mCherry (‘mACh’) or mCherry (‘mCh’) tagging to TPX2. PCR amplified a 3.O-kb fragment in the tagged line. Immunoblotting results are shown on the right; the band is shifted upwards with the tag. (F) PCR and immunoblotting confirmation of AKAP450 KO cells. DNA amplification was not observed when a primer targeting an exon was used for the KO line, whereas the hygromycin cassette was amplified with the primers designed at UTRs (this primer set did not amplify very long AKAP450 genes in the parental line). Immunoblotting showed a specific >250 kD band only in the parental line. (G) PCR and immunoblotting confirmation of CAMSAP3 KO. DNA amplification was not observed when a primer targeting an exon was used for the KO line. Immunoblotting showed a specific −150 kD band only in the parental line.

**Figure S2.**
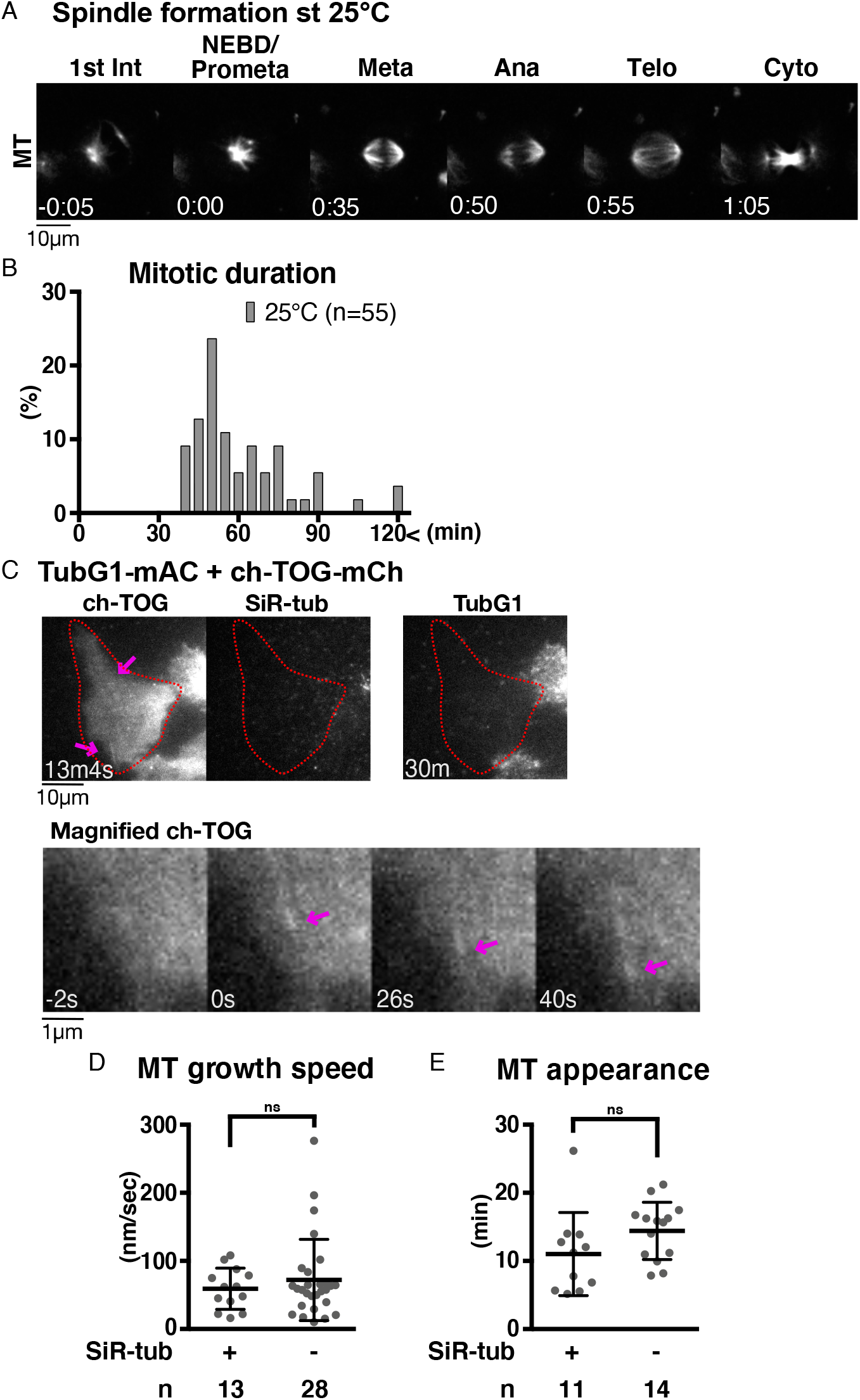
Validation of MT regrowth assay. (A) Mitosis of HCT116 cell line at 25 °C. Time 0 corresponds to NEBD. (B) Mitotic duration at 25 °C (NEBD to anaphase onset). (C) MT nucleation in the absence of γ-tubulin, without SiR-tubulin staining, in interphase. The cells with undetectable levels of γ-tubulin are marked in red circles. MTs were visualised using ch-TOG-mCherry (arrows). (D, E) MT dynamics of γ-tubulin-depleted cells based on ch-TOG-mCherry signals with or without SiR-tubulin staining. MT growth rate was determined based on kymographs of ch-TOG-mCherry. Statistical evaluation was performed using unpaired t-test with Welch’s correction (p = 0.3644, 0.1315).

**Figure S3.**
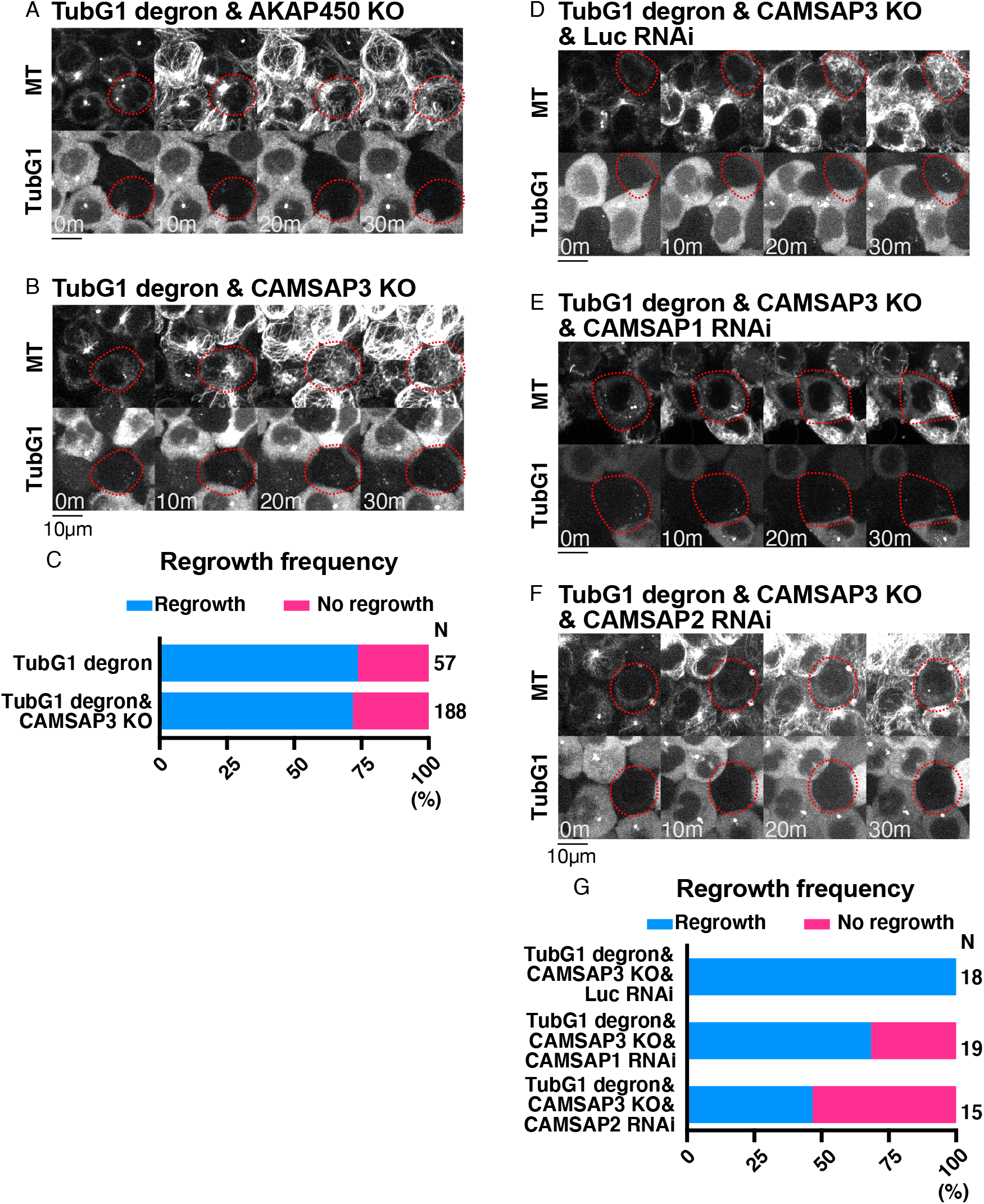
Additional data on MT regrowth ability in interphase cells. (A-B, D–F) MT regrowth in various lines. The cells marked in red circles show no detectable γ-tubulin signals. (C, G) Frequency of MT regrowth. MT appearance was assessed 30 min after nocodazole washout. The TubGl degron data in (C) are the duplicates of Fig. 6D.

**Figure S4.**
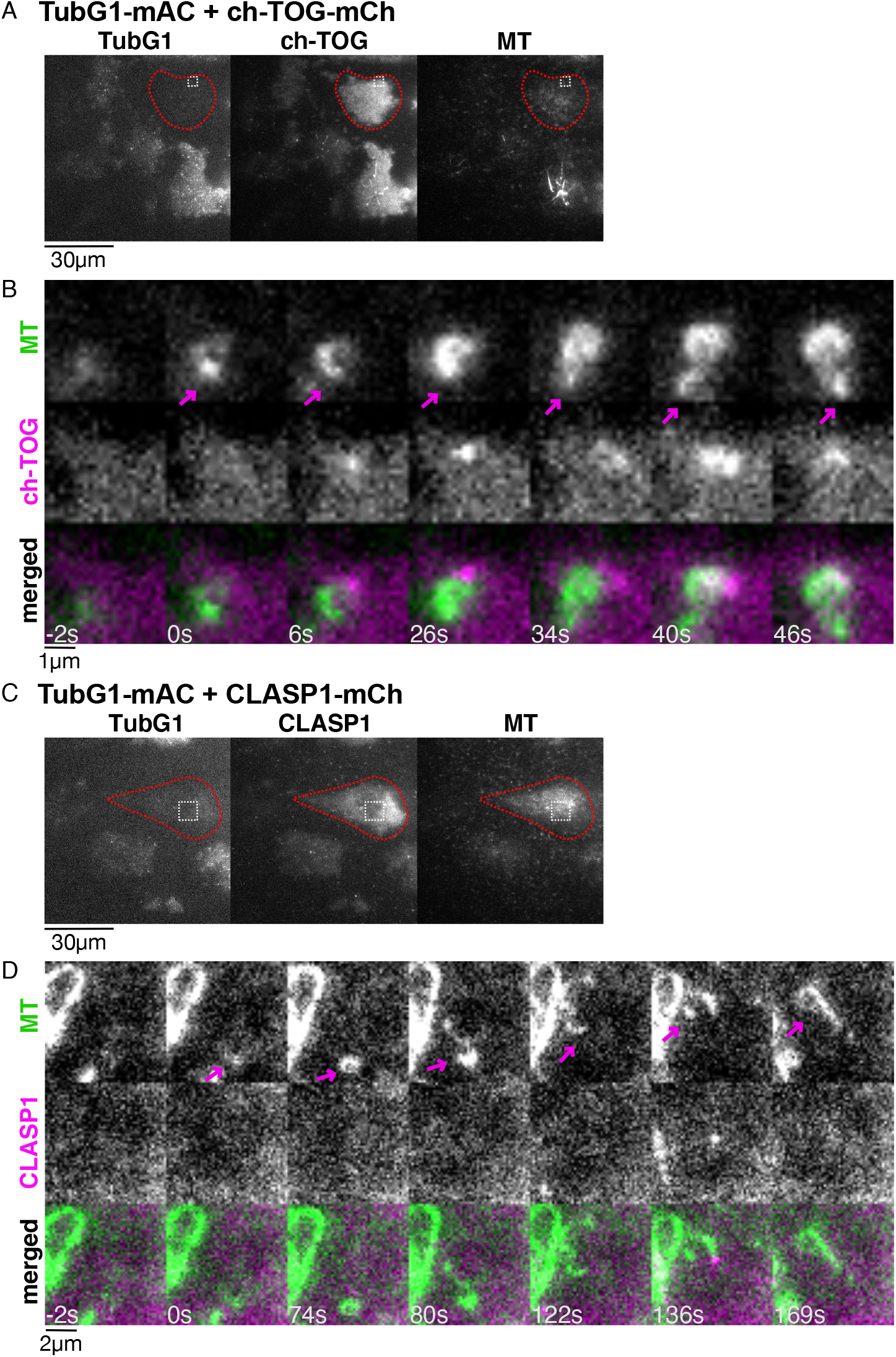
ch-TOG and CLASP1 localisation during γ-tubulin-independent MT nucleation in the interphase cytoplasm. Oblique illumination fluorescence microscopy of TubG1-mAID-mClover, ch-TOG-mCherry (or CLASPl-mCherry), and SiR-tubulin. Three-colour images were acquired to show γ-tubulin depletion in the first frame (A, C), followed by two-colour imaging every 2 s (B, D). A part of the cytoplasm (boxed in A, C) is magnified to show a nucleation event (B, D). Cells marked in red circles showed no detectable γ-tubulin signals.

**Figure S5.**
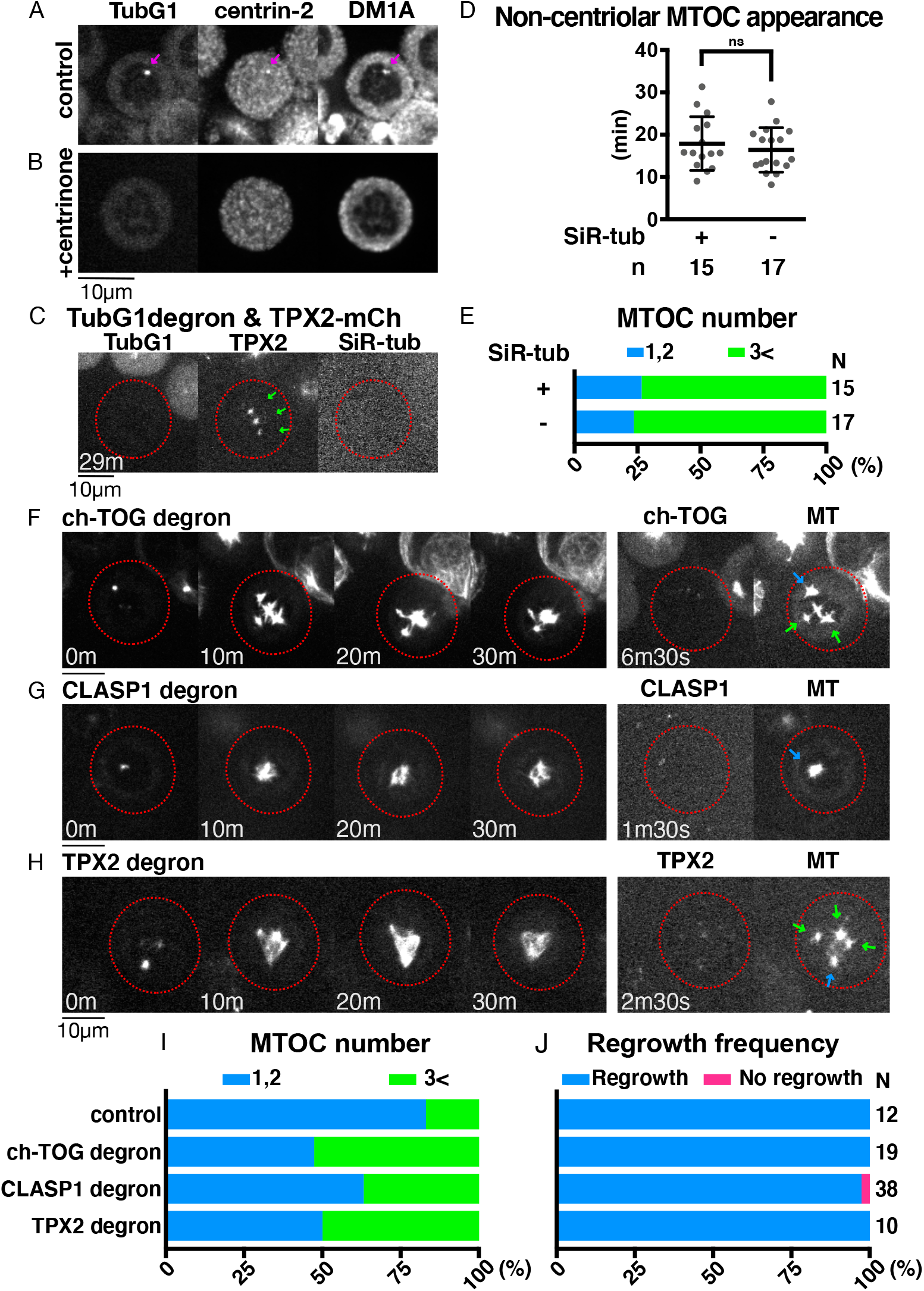
Additional data on MTOC formation and MT regrowth in mitosis. (A, B) Centriolar MTs remain even after rigorous MT depolymerisation with nocodazole (arrows). (C) MTOC formation in the absence of γ-tubulin, with or without SiR-tubulin in mitosis. The cells with undetectable levels of γ-tubulin are marked in red circles. Green arrows indicate non-centriolar MTOCs. TPX2-mCherry was used to visualise MTs. (D) The time of new MTOC appearance after nocodazole washout in the absence of γ-tubulin with or without SiR-tubulin staining (18 ± 6 min, 16 ± 5 min, p = 0.483 (unpaired t-test with Welch’s correction)). (E) Total MTOC numbers did not change with or without SiR-tubulin staining. (F-H) MT regrowth after ch-TOG, CLASP1, or TPX2 degron treatment. Red circles indicate the cells with undetectable levels of proteins. Note that γ-tubulin is intact in these lines. Bars, 10 μm. (I, J) Total MTOC numbers per cell and the frequency of cells with MTs at 30 min after nocodazole washout.

